# Cyborg Organoids: Implantation of Nanoelectronics via Organogenesis for Tissue-Wide Electrophysiology

**DOI:** 10.1101/697664

**Authors:** Qiang Li, Kewang Nan, Paul Le Floch, Zuwan Lin, Hao Sheng, Jia Liu

**Author notes:** These authors contributed equally (Qiang Li, Kewang Nan, Paul Le Floch).

## Abstract

Tissue-wide electrophysiology with single-cell and millisecond spatiotemporal resolution is critical for heart and brain studies, yet issues arise from invasive, localized implantation of electronics that destructs the well-connected cellular networks within matured organs. Here, we report the creation of cyborg organoids: the three-dimensional (3D) assembly of soft, stretchable mesh nanoelectronics across the entire organoid by cell-cell attraction forces from 2D-to-3D tissue reconfiguration during organogenesis. We demonstrate that stretchable mesh nanoelectronics can grow into and migrate with the initial 2D cell layers to form the 3D structure with minimal interruptions to tissue growth and differentiation. The intimate contact of nanoelectronics to cells enables us to chronically and systematically observe the evolution, propagation and synchronization of the bursting dynamics in human cardiac organoids through their entire organogenesis.

---

Complex organs (e.g. heart and brain) rely on spatiotemporally orchestrated communication of heterogenous cells to generate whole-organ functions.^1,2^ Thus, understanding development, mechanism and diseases of these organs requires system-level mapping of the cellular activities with high spatiotemporal resolutions across their entire 3D volumes over a substantial time window.^3–5^ Recently, tremendous progresses have miniaturized state-of-the-art electrophysiological tools into micro/nanoscales^6, 7^ with soft and multiplexed electronic units,^8, 9^ which have significantly reduced tissue disruptions while maintaining the unmatched spatiotemporal resolution for recording and manipulating.^10–13^ However, most of these devices either contact organs at the surface or penetrate locally and invasively through micro-needle injections. It remains a key challenge to uniformly implant and distribute a large number of interconnected and individually addressable sensors/stimulators throughout the 3D organs with minimal damage for chronic recording.

It is noteworthy that 3D organs originate from 2D embryonic germ layers during in vivo organogenesis. For instance, both neural and heart tubes originate from 2D cell layers in embryo via a series of cell proliferation and folding process and finally develop into brain, spinal cord and heart.^14, 15^ Such a 2D-to-3D transition in in vivo organogenesis has recently been reproduced in vitro for human organoids culture,^16, 17^ in which cell condensation from human mesenchymal stem cells (hMSCs) drives human induced pluripotent stems cells (hiPSCs) to fold into 3D organoids for transplantations and drug screening.^18, 19^

Inspired by the natural organogenesis, we propose a new scheme to realize 3D implantation and distribution of nanoelectronics within organoids (Figure. 1A). The first stage consists of transferring and laminating a mesh-like planar nanoelectronics together with its input/output (I/O) interconnects, onto a 2D sheet of stem cells (Stage I). Cell attraction forces between stem cells during cell aggregation, proliferation and migration gradually shrink the cell-sheet into a cell-dense plate, which simultaneously compresses the nanoelectronics into a closely packed architecture and covers the nanoelectronics with stem cells (Stage II). This interwoven cell/nanoelectronic structure then contracts and curls as a result of organogenesis-induced self-folding, first into a bowl geometry (Stage III), and then into a 3D spherical morphology (Stage IV). During this process, the mesh device seamlessly reconfigures with the cell-plate due to its soft mechanics, while maintaining uniform spatial distributions throughout the tissue, leading to a fully-grown 3D organoid with an embedded sensors/stimulators array in a minimally invasive and globally distributed manner (Stage IV) -- hence the name “cyborg organoid”. Finally, the stem cells in the as-formed cyborg organoid can further differentiate into targeted types of functional cells such as cardiomyocytes, while their electrophysiological activities can be chronically monitored using the embedded nanoelectronics (Stage V).

**Figure 1.**
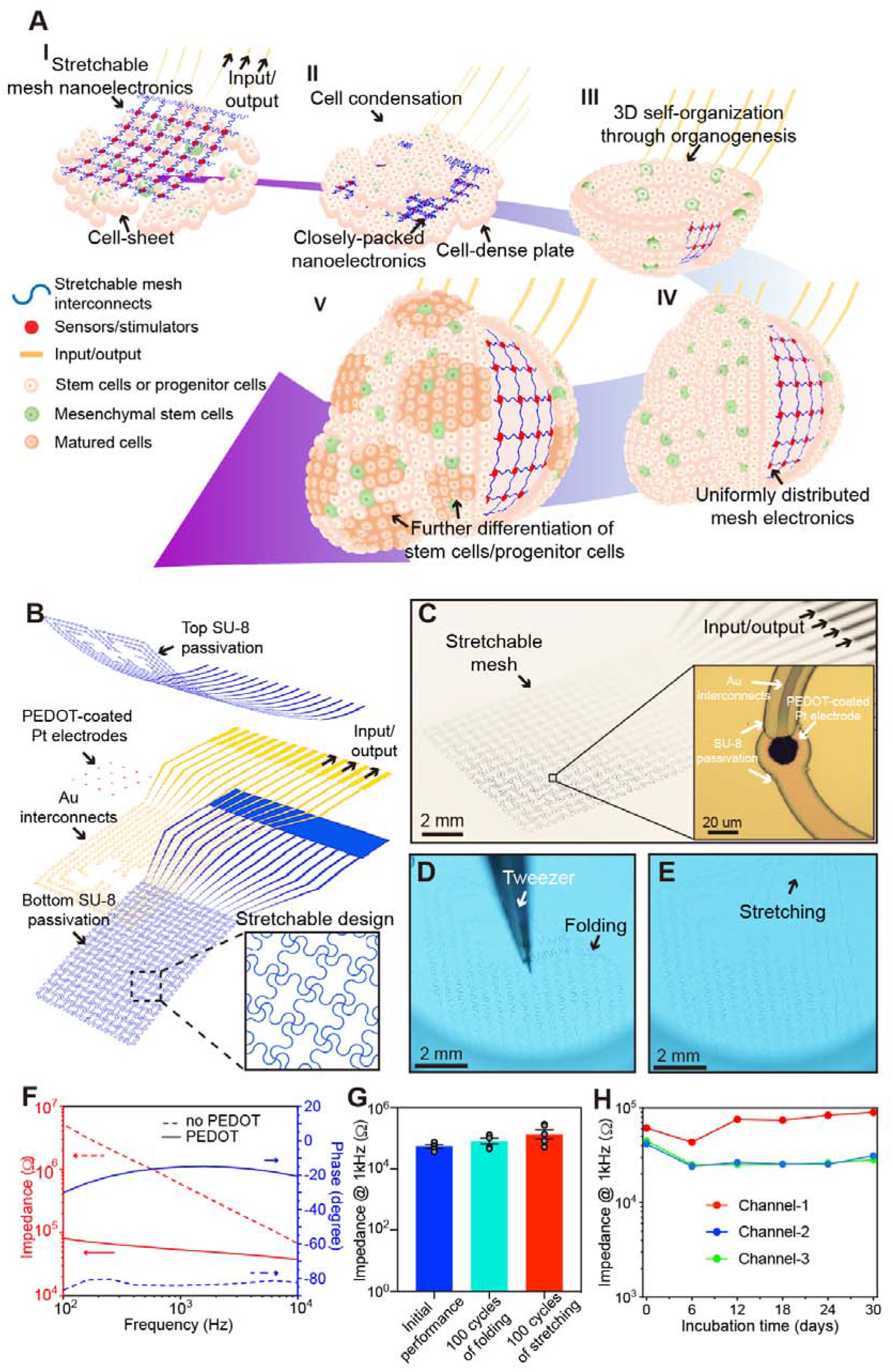
Cyborg organoids: concept and design. (A) Schematics illustrate the stepwise assembly of stretchable mesh electronics into organoids through organogenesis. (I) lamination of stretchable mesh electronics onto a continuous sheet of Matrigel seeded with Human induced pluripotent stem cells (hiPSCs) or hiPSCs-derived progenitor cells. (II) Aggregation of cell-sheet into a cell-dense plate through cell proliferations and migrations induced by cell-cell attractive forces, which embeds stretchable mesh electronics into the cell-plate and folds it into a closely packed structure. (III) Organogenetic 2D-to-3D self-organization folds 2D cell-plate/nanoelectronics hybrid into 3D structure. (IV) Organogenesis unfolds the closely packed nanoelectronics and distributes its structure across the entire 3D organoid. (V) Further development and differentiation of stem/progenitor cells into different cell types with their electrophysiological behaviors continuously monitored by 3D embedded nanoelectronics. (B) Exploded view of stretchable mesh nanoelectronics design, consisting of (from top to bottom) a 400 nm-thick top SU-8 encapsulation layer, a 50 nm-thick platinum (Pt) electrode layer coated with poly(3,4-ethylenedioxythiophene) (PEDOT), a 40 nm-thick gold (Au) interconnects layer, and a 400 nm-thick bottom SU-8 encapsulation layer. The inset shows the serpentine layout of the mesh. (C) Optical image of mesh electronics before releasing from the fabrication substrate. The inset shows the zoom-in view of a single Pt electrode coated with PEDOT. (D-E) Optical images of released stretchable mesh electronics in water, bent (D) and stretched (E) by tweezer. (F) Impedance and phase from 0.1 to 10 kHz of representative channels with and without PEDOT. (G) Average impedance (n=5) at 1 kHz for each channel in its original state, after 100 cycles of folding (to the degree as shown in (D)) and 100 cycles of stretching (to ca. 30% biaxially as shown in (E)). (H) Impedance at 1 kHz as a function of incubation time in PBS at 37°C for three representative channels.

We underline several important design characteristics of the nanoelectronics that enable the as-described seamless, whole-tissue-wide integrations (Figure. 1B). First, the device exploits a serpentine mesh layout with an overall filling ratio of less than 11%, which leads to significantly 20 improved in-plane stretchability up to 30% and compressibility up to several times of its initial volume due to out-of-plane buckling of the mesh network,^21^ thus capable of accommodating drastic volumetric changes (mostly compressive) during organogenesis.^16–19^ Second, two sets of mesh dimensions are implemented (ribbon width/thickness = 20 μm/0.8 μm or 10 μm/2.8 μm), which result in a tiny device mass of less than 15 μg and effective bending stiffness of 0.090 n·Nm and 1.9 n·Nm respectively (see Note S1 and Figure. S1 for details) that are essentially imperceptible to the surrounding tissues. Third, sensors with low-impedance and an average diameter of 20 μm that approaches the typical size of an individual cell^22^ enable non-invasive, localized, single-cell electrophysiological recordings. Detailed fabrication procedures can be found in the Supplementary Methods and Figure. S2.

Figure 1C shows mesh nanoelectronics immediately before release from the fabrication substrate, while the inset shows an individual platinum electrode electrochemically deposited with poly(3,4-ethylenedioxythiophene) (PEDOT) to further lower the interfacial impedance.^23^ Folding (up to 180°) and stretching (up to 30% biaxially) the released device in a chamber filled with water (Figure. S3) for 100 cycles (Figure. 1D-F and Supplementary videos 1-2) reveal neither visualizable damages (Figure. 1D-E) nor significant changes in the impedance of electrodes (Figure. 1F). Impedance measurements (Figure. S4) also show an >10 times reduction in impedance at 1 kHz and a shift of phase towards resistance-dominated behaviors after coating the electrodes with PEDOT.^23^ Furthermore, we test longevity of the device using a soaking test in 1x phosphate-buffered saline (PBS) at 37 °C, where impedance values of three random channels at 1 kHz remain steady over 30 days (Figure. 1H), showing the potential of such sensory platforms for chronic studies in physiological environments.

We integrate stretchable mesh nanoelectronics with hMSCs co-cultured hiPSC (Figures. S5-S12) or hiPSC-derived cardiac progenitor cells (hiPSC-CPCs) (Figure. 2 and Figures. S13-14). The initial flat cell-sheet sandwiching the nanoelectronics with Matrigel (Figure. 2A, I) shrinks into a cell-plate via hMSCs-driven condensation, spontaneously packaging the nanoelectronics into a highly compressed structure (Figure. 2A, II). Further 3D re-organization transforms the cell-plate into a 3D organoid (Figure. 2A, III) in ~72 hours. Phase images clearly illustrate initial compression (Figure. 2C, I to II) and subsequent folding (Figure. 2C, III) of the nanoelectronics together with cell-sheet/plate at each step. With further growth and expansion, the cyborg organoid fully unfolds and distributes the nanoelectronics throughout the entire organoid. Phase and false color images after ~480 hours of organogenesis clearly show uniform distribution of the device over and within the entire organoid as it grows and expands (Figure. 2D). Importantly, the same morphology from hMSCs/hiPSCs and hMSCs/hiPSC-CPCs assembled nanoelectronics further demonstrate the generality of this method for different types of cyborg organoids (Figure. S10 and Figure. 2A-C; Figure. S11 and Figure. S14).

**Figure 2.**
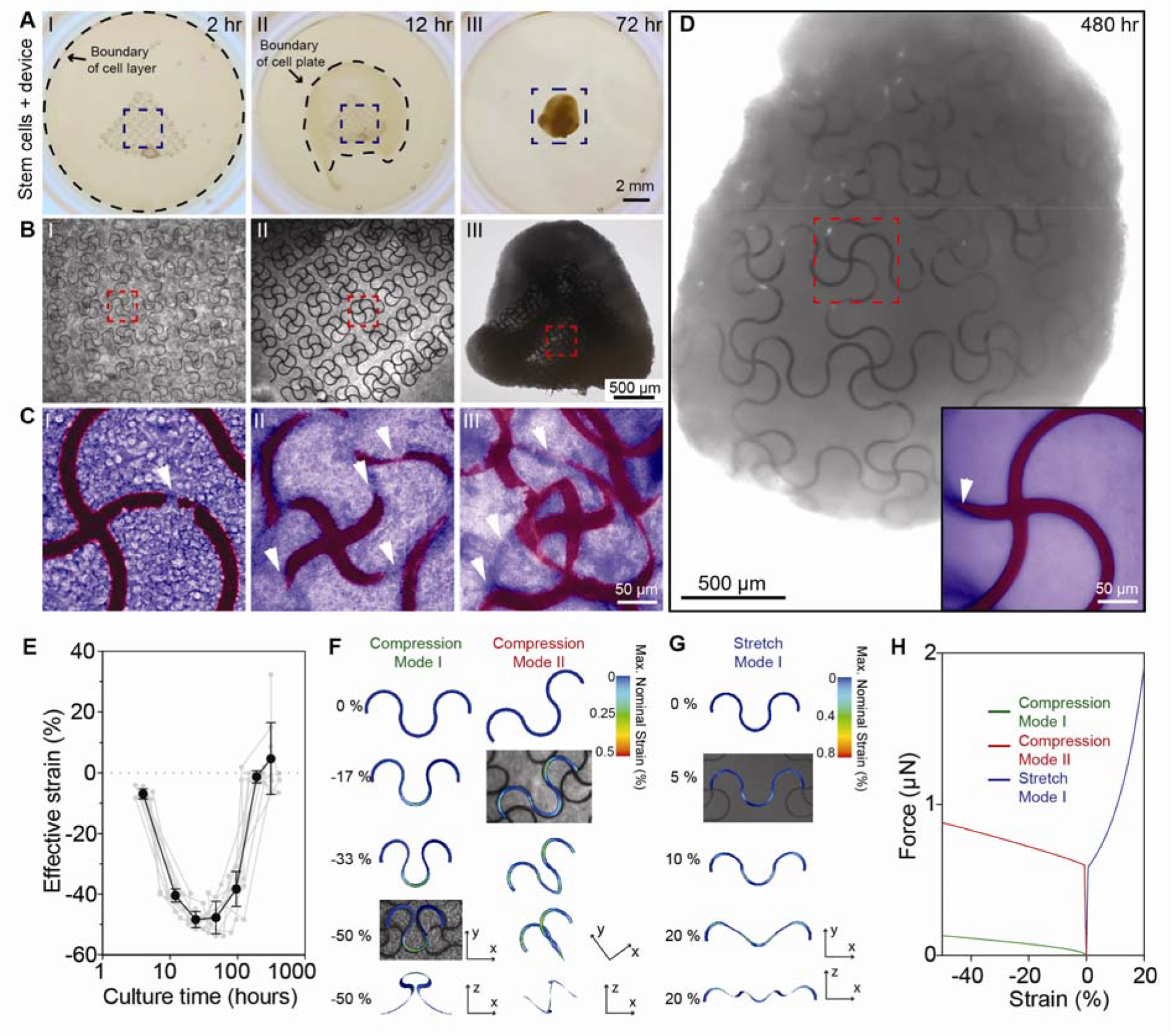
3D implantation and distribution of stretchable mesh nanoelectronics by organogenesis. (A) Optical images show representative steps of assembly of stretchable mesh nanoelectronics by organoids development corresponding to schematics in Figure. 1A, including lamination of stretchable mesh nanoelectronics on the stem cell cell-sheet on Matrigel (I), cell-dense cell-plate (II) and 3D re-organized cell-plate (III). hiPSCs-derived cardiac progenitor cells (hiPSCs-CPCs) and human mesenchymal stem cells (hMSCs) are cocultured. Dashed-line circles highlight boundaries of cell-plates at different stages. (B) Zoom-in phase images show the deformation of stretchable mesh nanoelectronics by the mechanical forces from organogenesis at different steps from the blue-dashed-box highlighted regions in (A). (C) Further zoom-in phase images with false color highlight the cell-nanoelectronics interaction at different steps from the red-dashed-box highlighted regions in (B). Red and blue false color label the SU-8 ribbons from nanoelectronics and stem cells, respectively. White arrows highlight the regions where the ribbons are embedded within the cells. (D) Bright-field phase image of a representative fully assembled cyborg organoid with stretchable mesh nanoelectronics unfolded and integrated across the entire organoid. Inset shows zoom-in phase image with false color from the red dashed box highlight region with the same color code as (C). (E) Representative effective strains of the device calculated through phase images as a function of culture time. Gray dots and lines show the individual data points, while black dots and lines show the averaged results. (Value = mean ± s.e.m., n=10). (F) Two compression modes of a repetitive unit in the mesh electronics obtained from finite element analysis (FEA). Merged simulation and optical images show the representative modes in real samples. The color scale indicates nominal strain in the unit. (G) A stretch mode of a repetitive unit in the mesh electronics obtained from FEA. Merged simulation and optical images show the representative modes in real samples. The color scale indicates nominal strain in the unit. (H) Magnitude of force as a function of strain for the three deformation modes in (F) and (G) obtained from FEA.

Imaging results show that during organogenesis, up to ~50% compressive and ~20% tensile effective strains, defined as the relative changes in distance between adjacent nodal points of the mesh, are applied to the nanoelectronics. Simulations by finite element analysis (FEA, see Note S2 for more details) show that the large deformations are accommodated by out-of-plane buckling of the mesh ribbons, leading to a maximum nominal strain under 1.2% which is below the fracture limit of both SU-8 and gold (Figure. 2F-G and Figures. S15-S19), indicating the robustness of this nanoelectronic structure design. Notably, buckling modes in compression and stretching observed in experiments can be qualitatively reproduced by the simulations. In addition, simulations show that the end-to-end force required to deform nanoelectronics is comparable to the forces required to deform cells at the same scale,^24^ indeed indicating a minimally invasive implantation process (Figure. 2H and Figure. S20).

At day 40 of tissue development and differentiation, optical images show that the nanoelectronics have been completely implanted into organoids, covering the entire tissue area with unfolded mesh ribbons (Figure. 3A). Flexibility and stretchability of the completely embedded nanoelectronics allow them to adapt to the contraction of the cardiac organoids (Supplementary videos 3-6). To further reveal the embedding of nanoelectronics within the 3D organoid, cyborg cardiac organoids at day 40 of differentiation are cleared. 3D reconstructed fluorescence images show the 3D interwoven nanoelectronics/cellular structures (Figure. 3B and Figure. S21) across the entire organoids. Zoom-in views further reveal that the nanoelectronics are sandwiched between the outer layer of the organoid consisting of migrated hMSCs (Figure. 3C, I) and the core of the organoid consisting of matured cardiomyocytes. Importantly, as planar nanoelectronics develop together with the 2D cell-plate, images show an intimate coupling between differentiated cardiomyocytes with nanoelectronics (Figure. 3C and Figure. S21B, D) throughout the entire organoid, which is rarely observed from other implantation methods for the post-matured living organoids and organs.

**Figure 3.**
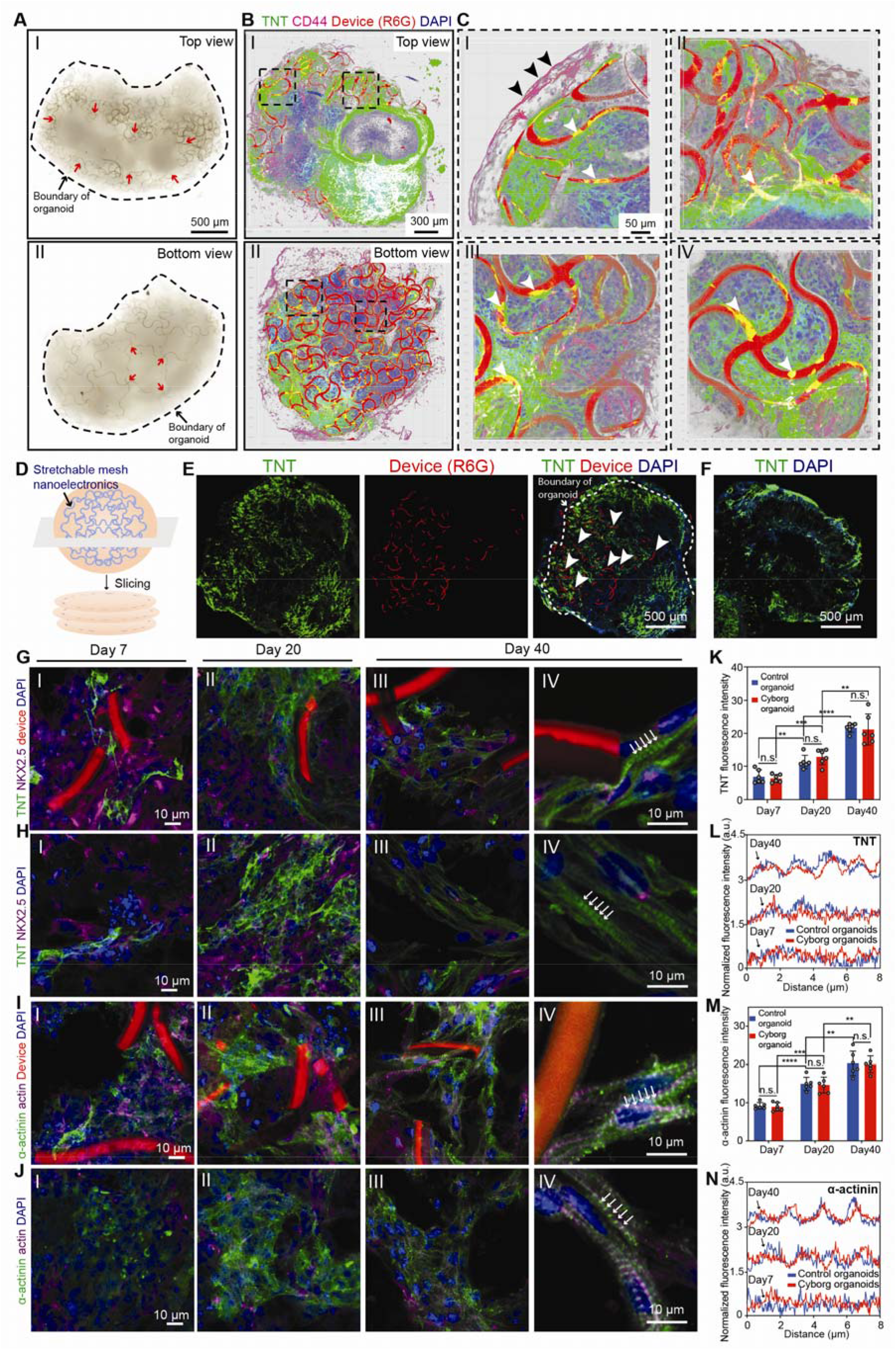
Development of human cardiac cyborg organoids. (A) Top (I) and bottom (II) views of the cardiac cyborg organoid at day 40 of differentiation. Red arrows highlight folding and stretching of stretchable mesh device by 3D re-organization of the cell-plate. (B) Top (I) and bottom (II) views of the 3D reconstructed fluorescence imaging of a cleared, immunostained cardiac cyborg organoid at day 40 of differentiation. Green, magenta and blue colors correspond to TNT, CD44 and DAPI, respectively. SU-8 in device was labeled by R6G for imaging in this and the following panels. (C) Zoom-in views from the dark dashed boxes highlighted region in (B) show that migrated hMSCs forming the outer layer of the organoid (I) and intimate coupling between iPSCs-derived cardiomyocytes with nanoelectronics (white arrows) were observed (IIV). (D) Schematics of slicing cardiac cyborg organoids for characterization at different stages of organogenesis. (E-F) Projection of 3D reconstructed confocal microscopic fluorescence images of the 30-μm-thick cyborg organoid (E) and control organoid (F). Green, red and blue colors correspond to TNT, device and DAPI, respectively, and are denoted at the top of the image panel in this and subsequent images. (G-N), Immunostaining and fluorescence imaging characterize the maturation of cardiomyocyte in the organoid at day 7 (I), 20 (II) and 40 (III-IV) of differentiation. TNT and NKX2.5 (G-H), and α-actinin and actin (I-J) were used to characterize the level of maturation of organoids with and without nanoelectronics embedding. Zoom-in views (IV) further show subcellular localization of TNT and α-actinin at day 40 of differentiation. Averaged fluorescence intensity of TNT (K) and α-actinin (M) over culture time. (Value = mean ± s.e.m., n= 6. **, P < 0.01; ***, P < 0.001; ***, P < 0.0001, two-tailed, unpaired t-test.). Fluorescence intensity plot showed the sarcomere alignment (L) and α-actinin alignment (N) in control and cyborg organoids at day 7, 20 and 40 of differentiation.

To investigate the effect of integrated nanoelectronics on the differentiation of organoids, hiPSC-CPCs/hMSCs cardiac cyborg organoids at different stages (day 7, 20 and 40 of differentiation) are fixed, sectioned, and immunostained for stage-specific marker expressions (Figure. 3D-F and Figures. S22-23), which are compared with those from control cardiac organoids. Protein markers including TNT, α-actinin and actin are imaged to quantify the maturation of cardiomyocytes while NKX2.5 is imaged for hiPSC-CPCs (Figure. 3G-J).^25^ Averaged fluorescence intensity over culture time shows significantly increased fluorescence intensity of TNT and α-actinin (Figure. 3K, M), while no statistically significant difference among cyborg and control organoids at all stages can be found. In addition, the fluorescence intensity plot along the axis of individual cells within the organoids shows the same level of increased sarcomere alignments in cyborg and control organoids from day 7 to day 40 of differentiation (Figure. 3L, N). These results substantially demonstrate that integrated nanoelectronics do not interrupt the 3D differentiation of iPSC-derived cardiac organoids.

The strong coupling between stem/progenitor cells and nanoelectronics offers us a unique opportunity to map the evolution of cellular electrophysiology during organogenesis. As a proof-of-concept, electrophysiological recordings (Figure. S24) are performed on cardiac cyborg organoids, which are typically formed in ~48 hours (Figure. 4A-B). Figure 4C shows the voltage trace of 14-channel (out of 16 channels) electrophysiological recording on cardiac cyborg organoid at day 35 of differentiation. The zoom-in plot of single spikes (Figure. 4D) show nonuniform electrophysiological behaviors of the cells distributed across the organoid, as well as a clear time latency revealing tissue-wide propagation of local field potentials (LFP). Chronic tracing of LFP at millisecond temporal resolution during organogenesis (Figure. 4E) reveals changes in the spike dynamics from initially slow waveform through the emergence of repolarization to fast depolarization.^26^ The averaged amplitude of the fast component associated with depolarization remains undetectable until day 31 of differentiation, and then increases monotonically (Figure. 4F). The field potential duration (FPD) is found to remain relatively steady between 0.7 and 0.8 s (Figure. 4G) with a slight increase. The activation mapping confirms the gradually synchronized LFP propagation across the cardiac organoid from day 26 to day 35 of differentiation (Figure. 4H and Figure. S25). These recording data suggest that the functional maturation of a cardiac organoid is marked by the synchronization of the bursting phase, instead of bursting frequency or FPD.

**Figure 4.**
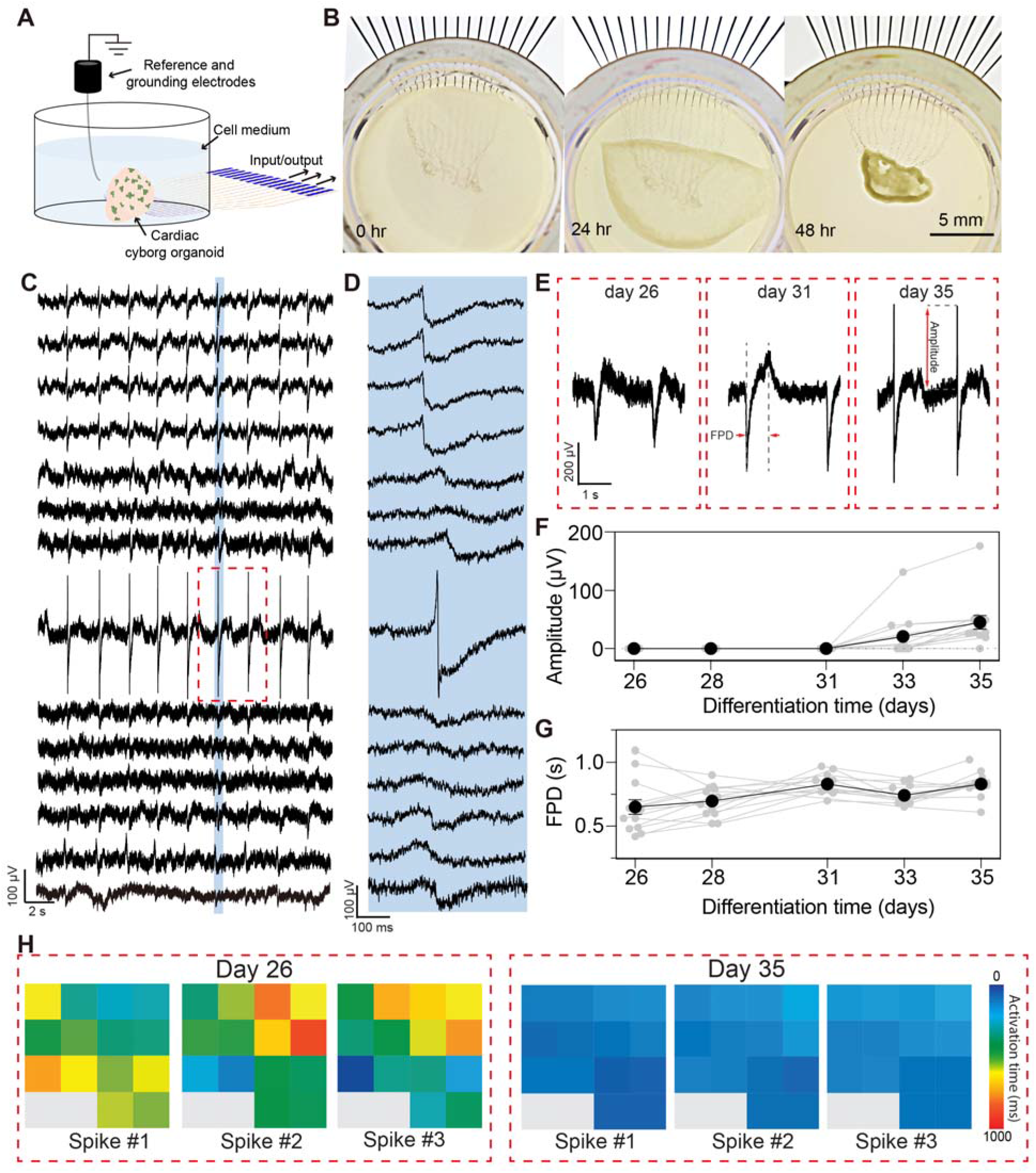
Chronic, multiplex, tissue-wide electrophysiological mapping of cardiac organoid during organogenesis. (A) Schematics show the set-up that connects cyborg cardiac organoids to external recording equipment for multiplexing electrophysiology during organoid development. (B) Optical images show representative processes (0, 24 and 48 hours) of 3D organization of cardiac cyborg organoids in culture chamber connected for electrophysiological recording. (C) 14-channel voltage traces recorded from the cardiac cyborg organoid at day 35 of differentiation. (D) Zoom-in views of the blue box highlights in (C) show single-spiked field potential recording. (E) Zoom-in views of red dashed box highlighted channel in (C) on three different culturing days (day 26, 31, and 35 of differentiation). (F-H) Amplitude of fast peak (F) and field potential duration (G) defined in (E) as a function of differentiation time. Gray lines show individual channels. Black line shows the averaged results from 14 channels. (Value = mean ± s.e.m., n= 14). (H) Isochronal mappings at day 26 and day 35 of differentiation show delays of activation time (max. -dV/dt) for three consecutive spiking events labeled in Figure. S25.

In summary, we have created the first human cardiac cyborg organoid via organogenetic 2D-to-3D tissue reconfiguration and studied the evolution of electrophysiological patterns during organogenesis. This method is ready to be scalable for integrating a larger number of sensors fabricated by photo- or electron beam-lithography.^27^ Additional work remains to apply cardiac cyborg organoids to study cardiac development, diseases and therapeutics.^28^ Further development and generalization of cyborg organoids could serve as a paradigm-shifting platform for spatially resolved, high-fidelity and chronic electrophysiological recordings for many other types of organoids^29, 30^ and animal embryos as well as for monitoring and controlling of organoid-enabled cellular therapeutics.^31^

## AUTHOR INFORMATION

Corresponding Author

Author Contributions

*These authors contributed equally.

Notes

The authors declare no competing financial interest.

## ACKNOWLEDGMENT

We are grateful for the support of all Liu group members. We thank D. Mélançon for helpful discussions on FEA simulation, and T. Ye and X. Dai for helpful discussions on the electrophysiological recording. We also thank Prof. C. M. Lieber, Prof. Z. Suo, J. C. Salant and D. R. Lilienthal for suggestions on the manuscript. This research was partly supported by the Harvard Dean’s Fund for Promising Scholarship and Harvard University Center for Nanoscale Systems supported by NSF.

