## Supplementary document for "Cyborg Organoids: Implantation of Nanoelectronics via Organogenesis for Tissue-Wide Electrophysiology"

*These authors contributed equally.

**This PDF file includes:**

Materials and Methods

Supplementary Notes 1-2

Figures S1-S25

**Other Supplementary Materials for this manuscript include the following:**

Movies S1 to S6

Materials and Methods

1. **Fabrications of soft, stretchable mesh nanoelectronics**

Fabrications of the ultra-thin, stretchable mesh nanoelectronics were based methods described previously. ^1,2^ The key steps, illustrated in Figure. S2, included:

(**1**) Cleaning a silicon wafer grown with thermal oxide (500-nm thickness) with acetone, isopropyl alcohol, and water.

(**2**) Depositing 100-nm-thick nickel (Ni) using electron-beam evaporator as a sacrificial layer.

(**3**) Spin-coating SU-8 precursor (SU-8 2000.5, MicroChem) at 4000 rpm, which was pre-baked at (65 °C, 95 °C) for 2 min each, exposed to 365 nm ultra-violet (UV) for 200 mJ/cm^2^, post-baked at (65 °C, 95 °C) for 2 min each, developed using SU-8 developer (MicroChem) for 60 s, and hard-baked at 180 °C for 40 min to define mesh SU-8 patterns (400-nm thickness) for bottom encapsulation. For fluorescence imaging, 0.004 wt‰ of Rhodamin 6G powder (Sigma-Aldrich) was added into SU-8 precursor.

(**4**) Spin-coating LOR3A photoresist (MicroChem) at 4000 rpm, followed by pre-baking at 180 °C for 5 min; spin-coating S1805 photoresist (MicroChem) at 4000 rpm, followed by pre-backing at 115 °C for 1 min; the sample was then exposed to 405 nm UV for 40 mJ/cm^2^, and developed using CD-26 developer (Micropost) for 70 s to define interconnects patterns.

(**5**) Depositing 5/40/5-nm-thick chromium/gold/chromium (Cr/Au/Cr) by electron-beam evaporator, followed by a standard lift-off procedure in remover PG (MicroChem) overnight to define the Au interconnects.

(**6**) Repeating Step (**4**) to define electrode array patterns in LOR3A/S1805 bilayer photoresists;

(**7**) Depositing 5/50-nm-thick chromium/platinum (Cr/Pt) by electron-beam evaporator, followed by a standard lift-off procedure in remover PG (MicroChem) for 10 min to define the electrode array;

(**8**) Repeating Step (**3**) for top SU-8 encapsulation (400-nm thickness).

(**9**) Electrochemically polymerizing poly(3,4-ethylenedioxythiophene) (PEDOT) on the Pt electrode array using a precursor consisting of 14 mM 3,4-ethylenedioxythiophene (EDOT, Sigma-Aldrich) in 1x phosphate-buffered saline (PBS) solution. The precursor was drop-casted onto the device, followed by passage of a 1 V DC voltage (Dr. Meter DC power supply PS-305DM) for 200 s using device electrodes as the anode and an external Pt wire as the cathode. The device was then rinsed with DI water for 30 s and dried.

(**10**) Soldering a 16-channel flexible flat cable (Molex) onto the input/output pads using a flip-chip bonder (Finetech Fineplacer).

(**11**) Gluing a chamber onto the substrate wafer to completely enclose the mesh part of the device using a bio-compatible adhesive (Kwik-Sil, WPI).

(**12**) Treating surface of the device with light oxygen plasma (Anatech 106 oxygen plasma barrel asher), followed by adding 3 mL of Ni etchant (type TFB, Transene) into the chamber for 2 to 4 hours to completely release the mesh electronics from the substrate wafer. The device was then ready for subsequent sterilization steps before cell culture.

2. **Organoid culture**

**2.1 Materials for cell culture**

Human induced pluripotent stem cells hiPSCs-(IMR90)-1 were purchased from WiCell Research Institute (Madison, WI, USA). Authentication and test for the free of mycoplasma were performed by WiCell Research Institute. A cell bank was established after the cells were received. For experiments, each aliquot from the cell bank was used for less than 5 additional passages. Human mesenchymal stem cells (hMSCs; PT-2501) were obtained from Lonza (Walkersville, MD, USA) and used for less than 5 additional passages. Reagents and their supplies: Essential 8™ (E8) medium (cat # A1517001, life technologies); MSCGM BulletKit (cat # PT-3238 & PT-4105, Lonza); RPMI 1640 Medium (cat # 11875093, life technologies); B-27 Supplement (cat # 17504044, life technologies); B-27™ Supplement, minus insulin (cat # A1895601, life technologies); CHIR 99021, (cat # 44-231-0, Tocris Bioscience™); IWR-1 (cat # CAYM-13659-5, Cayman Chemical); Normal donkey serum (cat # 017-000-121, Jackson ImmunoResearch); Accutase (cat # A1110501, life technologies); Trypsin-EDTA 0.05% (cat # 25-300-054, Invitrogen); EDTA 0.5M pH 8.0 (cat # AM9260G, Invitrogen); Y-27632 (cat # 129830-38-2, Sigma); Matrigel (cat # 08-774-552, Corning); DAPI (cat # D9542, Sigma). Phalloidin-iFluor 647 (cat # ab176759, Abcam); Poly-D-lysine hydrobromide (cat # P7280-5MG, Sigma); Photoinitiator 2,20-Azobis[2-(2-imidazolin-2-yl) propane] dihydrochloride (cat # VA-044, Wako Chemicals). Antibodies used in this study: Nanog (Santa Cruz, 1:100); CD44 (Abcam, 1:100); Cardiac troponin T (TNT) (Invitrogen, 1:100); a-actinin (Sigma, 1:100); NKX2.5 (Abcam, 1:100); Alexa 594 donkey anti-mouse (Jackson ImmunoResearch, 1:500); Alexa 647 Donkey anti-rabbit (Invitrogen, 1:500).

**2.2 Two dimensional (2D) cells culture**

hiPSCs-(IMR90)-1 were maintained in 6-well plate coated with Matrigel in Essential 8^TM^ medium based on methods described previously.^3,4^ Steps were as follows:

(**1**) 6-well plate was coated with Matrigel at 37^o^C at least one hour.

(**2**) Matrigel from the well was aspirated and replaced with 2 mL E8 medium.

(**3**) hiPSCs were incubated with 0.5 mM EDTA at room temperature for 5 minutes.

(**4**) EDTA solution was aspirated and replaced with 1 mL E8 medium. Cells were pipetted into clusters.

(**5**) 100-200 µL cells were seeded in the Matrigel coated 6 well-plate.

(**6**) Medium was changed daily. Cells were passaged every 4 days.

hMSCs were cultured in T-75 flasks in MSCGM medium according to Lonza introduction manual. Steps were as follows:

(**1**) 5000-6000 cells per cm^2^ were seeded in T-75 flask with about 30 ml MSCGM medium.

(**2**) Medium was changed every 3-4 days.

(**3**) When hMSCs reached ~ 90% confluence, cells were dissociated with Trypsin-EDTA 0.05%. The cells were used by passage 5.

**2.3 2D hiPSC-derived cardiomyocytes differentiation**

Cardiomyocytes differentiation was based on methods described previously.^5,6^ Steps were as follows:

(**1**) hiPSCs were cultured in 6-well plate with E8 medium for 3-4 days.

(**2**) E8 medium was removed. RPMI 1640 medium plus 1% B27-insulin and 12 μM CHIR99021 were added for differentiation. This day was defined as Day 0.

(**3**) On day 1, the medium was changed to RPMI 1640 medium plus 1% B27-insulin.

(**4**) On day 3, the medium was changed to RPMI 1640 medium plus 1% B27-insulin and 5 μM IWR1.

(**5**) On day 5, the medium was changed to RPMI 1640 medium plus 1% B27-insulin.

(**6**) On day 7, the medium was changed to RPMI 1640 medium plus 1% B27.

(**7**) The medium was then replaced with fresh RPMI 1640 medium plus 1% B27 every other day. The cells started beating from day 8 or day 10.

**2.4 3D culture of cyborg organoids**

The integration of stretchable mesh nanoelectronics with cells to create cyborg organoids followed immediately from fabrications and release of the device in the chamber, which involved the following steps:

(**1**) The released device was sterilized by 1) DI water rinse for three times, and 2) 70% ethanol for at least 15 minutes.

(**2**) The device was washed with PBS three times followed by incubating with Poly-D-lysine hydrobromide (0.01% w/v) overnight.

(**3**) Poly-D-lysine hydrobromide solution was removed, and 300 μL liquid Matrigel (10 mg/mL) was added to the chamber from the device-free side on ice. The device was incubated for at least 30 minutes at 37°C to solidify the Matrigel layer.

(**4**) hiPSCs or hiPSC-derived cardiac progenitor cells (hiPSC-CPCs) (2~3×10^6^ cells) and hMSCs (2~4×10^5^ cells) were suspended in a mixture of E8 or RPMI 1640 (supplemented with 1% B27-insulin or 1% B27) and MSCGM medium, and then transferred onto the cured Matrigel in the chamber and cultured at 37°C, 5% CO_2_.

(**5**) For electrophysiological measurement, hiPSC-CPCs at day 24 and hMSCs mixture were transferred onto the Matrigel and cultured at 37°C, 5% CO_2_. hiPSCs-derived cyborg cardiac organoid was formed within 48-72 hours. Electrical activity from the organoid was recorded from 48 hours of its formation, which corresponded to day 26 of cardiac differentiation in Figure. 4.

3. **Characterization**
   3.1 **Organoids clearing, section, immunostaining and imaging**

For whole organoids imaging, procedures were adapted from tissue clearing techniques CLARITY^7^ and passive clarity technique (PACT),^8^ the organoids were fixed with 4% PFA at 4°C overnight and incubated with hydrogel solution (0.25% (w/v) VA-044 and 4% (w/v) acrylamide in PBS) at 4°C for 24 hours. The samples were placed in X-CLARITY hydrogel polymerization device for 3-4 hours at 37°C with -90 kPa vacuum, followed by wash in PBS overnight before electrophoretic lipid extraction for 24 hours in the X-CLARITY electrophoretic tissue clearing (ETC) chamber. Then immunostaining was performed by staining the primary antibodies for 3-5 days and the secondary antibodies for 2-4 days, respectively. The samples were submerged in optical clearing solution overnight and embedded in 2% agarose gel before imaging using Zeiss 2-photon time-lapse confocal microscopy at Harvard Center for Biological Imaging (HCBI). For characterization of cyborg cardiac organoids, the organoids at different stages were fixed with 4% PFA at 4 °C overnight and immersed in 30% sucrose for at least 12 hours. Then the samples were embedded in optimal cutting temperature (OCT) compound and cryostat section of 30 μm-thick slices. Cardiac organoids without device integrating were used as control. For staining, the first antibodies were incubated at 4 ^o^C overnight and the secondary antibodies were stained at RT for 3-4 hours. For 2D culture, cells were fixed with 4% paraformaldehyde (PFA) at room temperature (RT) for 15 minutes. Cells were incubated with primary antibodies at 4 °C overnight and the secondary antibodies were stained at RT for 1-2 hours. Finally, 4’,6-diamidino-2-phenylindole (DAPI) were stained for 10 minutes. Cells were washed with PBS three times before imaging. All samples were imaged by Zeiss 880 confocal microscopy at HCBI. Imaging was analysis by Zen (Blue edition) and Fiji. Fluorescence intensity was calculated by Fiji. Data analysis and statistical tests were performed by Graphpad Prism.

**3.2 Device impedance characterization**

A three-electrodes setup was used to measure the electrochemical impedance spectrum of the electrodes from each device. Platinum wire (300 μm in diameter, 1.5 cm in length immersed) and a standard silver/silver chloride electrode were used as counter electrode and reference electrode, respectively. The device was immersed in 1xPBS solution during measurement. The SP-150 potentiostat (Bio-logic) along with its commercial software EC-lab was used to perform the measurements. For each measurement, at least three frequency sweeps were measured from 1 MHz down to 1Hz to obtain statistical results. A sinusoidal voltage of 100 mV peak-to-peak was applied. For each data point, the response to 10 consecutive sinusoids (spaced out by 10% of the period duration) was accumulated and averaged. Optical images of the measurement setup can be found in Figure. S4.

4. **Electrophysiological measurement and data analysis**

The Blackrock CerePlex Direct voltage amplifier along with a 32 channels Blackrock μ digital headstage connected to the device was used to record electrical activity from organoids. The headstage-to-device connector (16 channels only) was homemade. The organoid culture media was grounded to earth and a reference electrode was also inserted in the media, far from the device (distance above 1 cm). Platinum wires were used as ground and reference electrodes. A sampling rate of 2000 samples per second was used. A 0.3 - 1000 Hz band-pass filter (Butterworth, 4th order) was applied to reduce high frequency noise. Matlab codes provided by Blackrock were used to convert raw data files into accessible format. Data were then transferred to Graphpad Prism for post-processing. Schematics of the measurement setup can be found in Figure. S24.

**Supplementary Notes**

**1. Bending stiffness calculations**

The bending stiffness of the serpentine mesh can be estimated using a beam model. For a homogeneous beam of elastic modulus *E*, width *w*, and thickness *h*, the bending stiffness is:

$$D=\frac{Ewh^{3}}{12}$$

$D=\frac{\mathrm{Ew}h^{3}}{12}$We derive the expression of the bending stiffness for a three-layer beam described in Fig. S1.

The bending moment along the x direction (per unit width) is:

$$m_{xx}=\int_{-h_{p}/2}^{h_{p}/2} z\sigma_{xx}dz$$

which can be separated into three terms:

$$m_{xx}=\int_{-h_{p}/2}^{-h_{m}/2} z\sigma_{xx}dz+\int_{-h_{m}/2}^{h_{m}/2} z\sigma_{xx}dz+\int_{h_{m}/2}^{h_{p}/2} z\sigma_{xx}dz$$

The normal stress along the x direction, σ_xx_, is given by Hooke’s law,

$$\sigma_{xx}=\frac{E}{1-\nu^{2}}(\varepsilon_{xx}+\nu\varepsilon_{yy})$$

In small deformations,

$$\varepsilon_{xx}=-z\frac{\partial^{2}w}{{\partial x}^{2}}$$

$$\varepsilon_{yy}=-z\frac{\partial^{2}w}{{\partial y}^{2}}$$

where w is the displacement in the z direction.

Thus,

$$z\sigma_{xx}=-z^{2}\frac{E}{1-\nu^{2}}(\frac{\partial^{2}w}{{\partial x}^{2}}+\frac{\partial^{2}w}{{\partial y}^{2}})$$

And the bending moment (per unit width) can be rewritten as:

$$m_{xx}=-\left( \int_{-h_{p}/2}^{-h_{m}/2} z^{2}\frac{E_{p}}{1-{\nu_{p}}^{2}}dz+\int_{-h_{m}/2}^{h_{m}/2} z^{2}\frac{E_{m}}{1-{\nu_{m}}^{2}}dz+\int_{h_{m}/2}^{h_{p}/2} z^{2}\frac{E_{p}}{1-{\nu_{p}}^{2}}dz \right)(\frac{\partial^{2}w}{{\partial x}^{2}}+\frac{\partial^{2}w}{{\partial y}^{2}})$$

where *E_p_* and *E_m_* are respectively the Young’s modulus of SU-8 and Gold, 2 and 79 GPa, and *ν_p_* and *ν_m_* are respectively the Poisson’s ratio of SU-8 and Gold, 0.22 and 0.43. The bending stiffness per unit width for the core-shell beam of widths *w_p_* and *w_m_* (geometry described in Fig. S1 is estimated as:

$$D=\frac{1}{12}\frac{E_{p}}{1-{\nu_{p}}^{2}}{(w}_{p}{h_{p}}^{3}-{{w_{m}h}_{m}}^{3})+\frac{1}{12}\frac{E_{m}}{1-{\nu_{m}}^{2}}{w_{m}h_{m}}^{3}$$

For the first mesh design (width/thickness = 20 μm/0.8 μm), where *h_p_* = 0.8 μm, *h_m_* = 0.04 μm, *w_p_* = 20 μm and *w_m_* = 10 μm, *D* = 1.8 10^-15^ N.m^2^. The flexural rigidity (or bending stiffness per unit width is):

$$d=\frac{D}{w_{p}}$$

And *d* = 0.090 nN.m.

Repeating for the second mesh design (width/thickness = 10 μm/2.8 μm), where *h_p_* = 2.8 μm, *h_m_* = 0.04 μm, *w_p_* = 10 μm and *w_m_* = 8 μm, *D* = 3.8 10^-14^ N.m^2^. Similarly, *d* = 1.9 nN.m.

**2. Numerical mechanical simulations**

Abaqus 6.12 was used to perform mechanical analysis of the structure. The goal of the simulations is to evaluate strain and stress concentration, as well as the effective stiffness of a unit ribbon while buckling. Figure 2E shows that the mesh undergoes a maximum compression of about 50% of its initial dimensions during the organoid development, as well as a maximum stretch up to 20%. Buckling of the ribbon structure is also observed experimentally.

We first analyze the buckling modes in compression of the unit ribbon using the “linear perturbation, Buckle” module of Abaqus for the two geometries of device (width/thickness = 10/2.8 or 20/0.8 μm). We show the first five modes of buckling in Fig. S15 and S16. One end of the ribbon is subjected to an encastre boundary condition while the second end is subjected to an in-plane displacement (rotation is constrained). As Mode I and Mode II in compression are the most widely observed in the optical images recorded, we only simulated these modes in large deformation. Similarly, we only simulated the Mode I during stretching of the device. To do so, a “static, general” step was run to apply an end-to-end compression of 50% or a stretch of 20%. Small concentrated forces are applied (magnitude ~10^-6^ - 10^-7^ μN) to induce preferred buckling modes. To check convergence, we varied the seeding size of the mesh (hexagonal elements) from 0.5 to 1 μm.

As expected, the strain is higher in the thicker device (10-μm width, 2.8-μm thickness). Still, the maximum principal nominal strain is below 1.2% in both the SU-8 and gold layers for the compression mode I (Fig. 2F and S17) and mode II (Fig. 2F and S18) and the stretching mode I (Fig. 2G and S19) that is well below the fracture limit of both materials. This result confirms that the device can resist both compression and stretch during organogenesis.

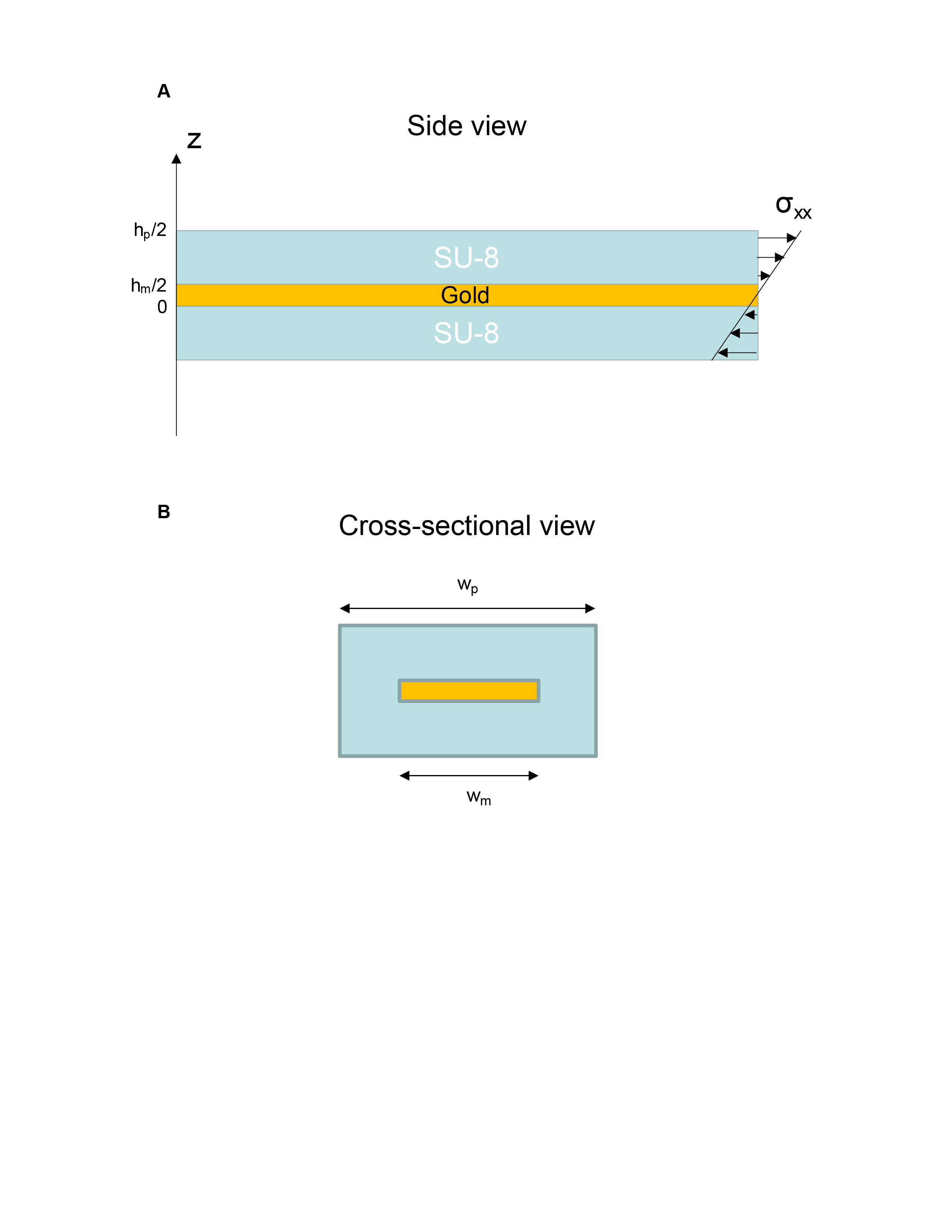

**Figure S1.** **Schematics show the model for effective bending stiffness calculation of the mesh nanoelectronics.** (A) The side view of individual ribbon on mesh nanoelectronics. (B) The cross-sectional view of individual ribbon on mesh nanoelectronics.

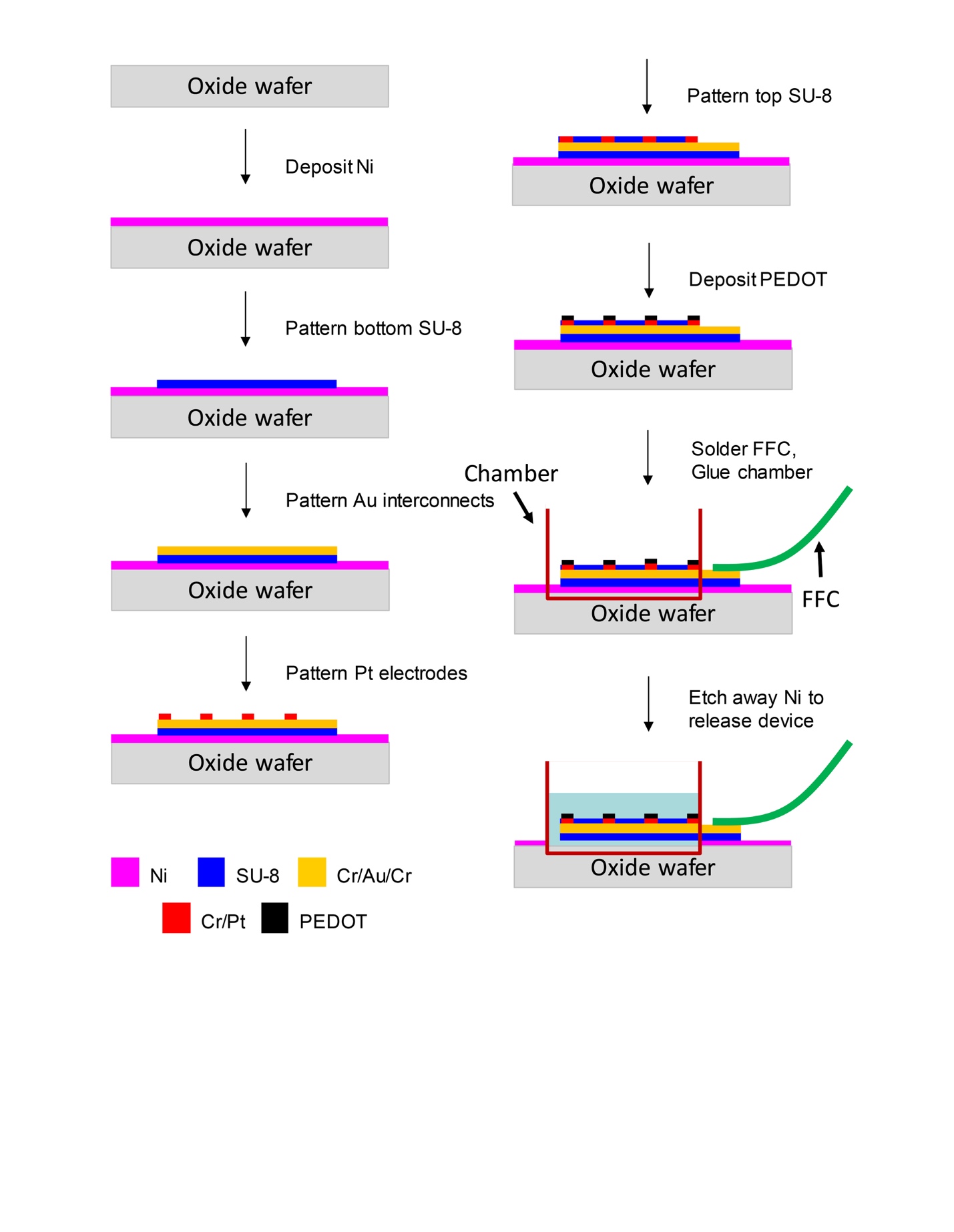

**Figure S2.** **Schematics show the entire fabrication flow of stretchable mesh nanoelectronics.**

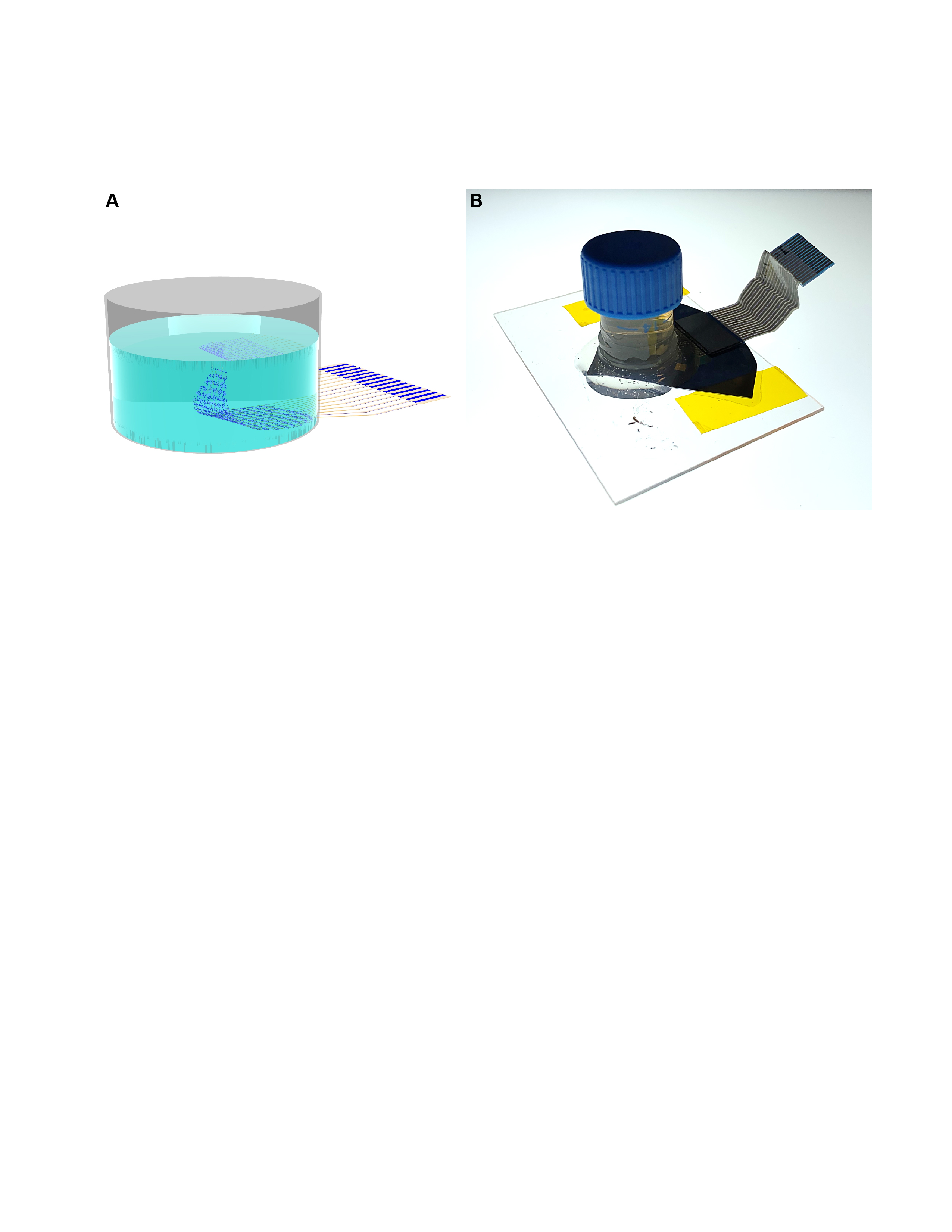

**Figure S3. A chamber is glued onto the device for releasing of mesh electronics in PBS solution.** (A) The schematic illustration. (B) The corresponding optical image.

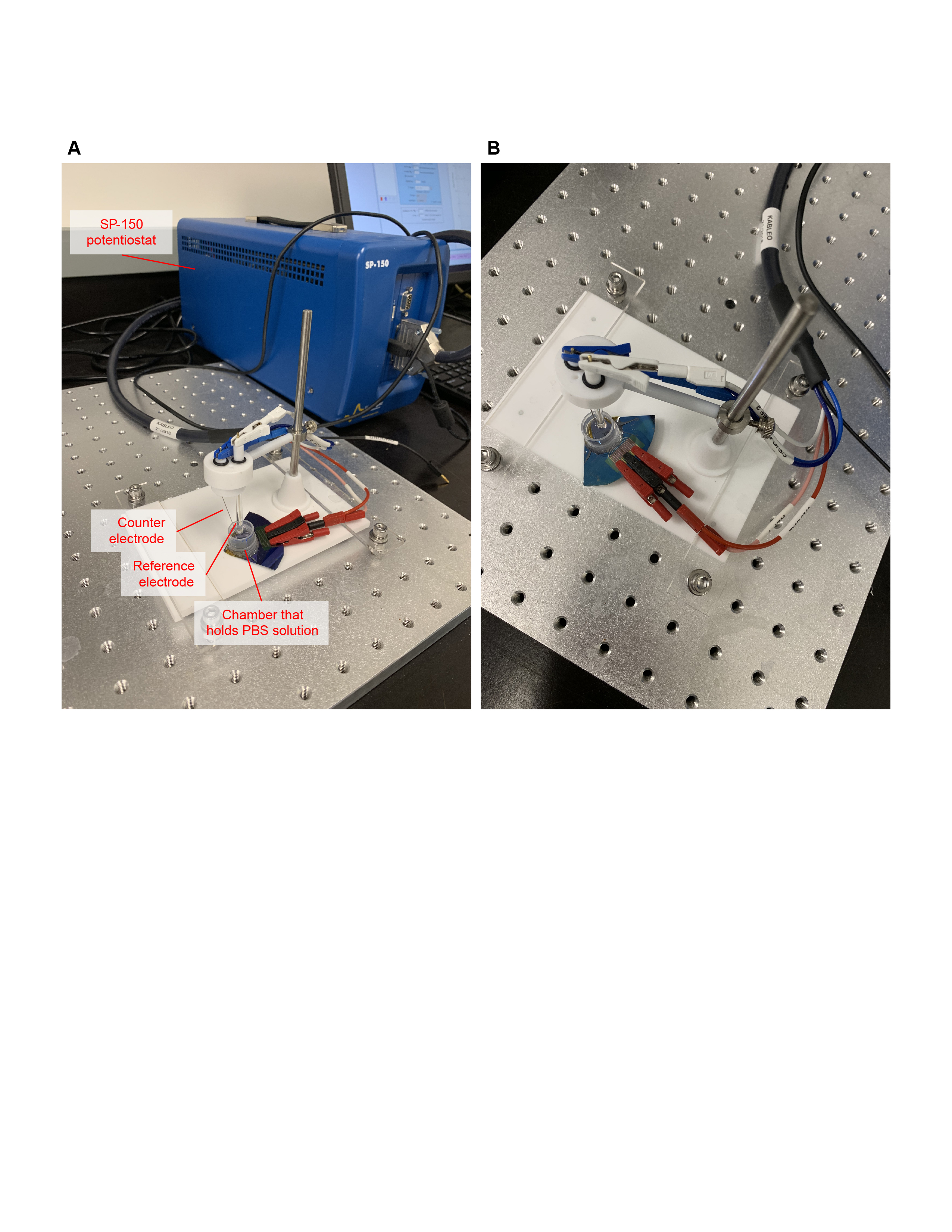

**Figure S4.** **Optical images show the electrochemical impedance spectrum (EIS) measurement setup for device characterization.** Reference and ground electrodes are directly inserted into the chamber containing the cyborg organoid in PBS solution. Crocodile clips are used to connect individual channels to the SP-150 potentiostat for EIS recording. (A) The perspective view. (B) The top view.

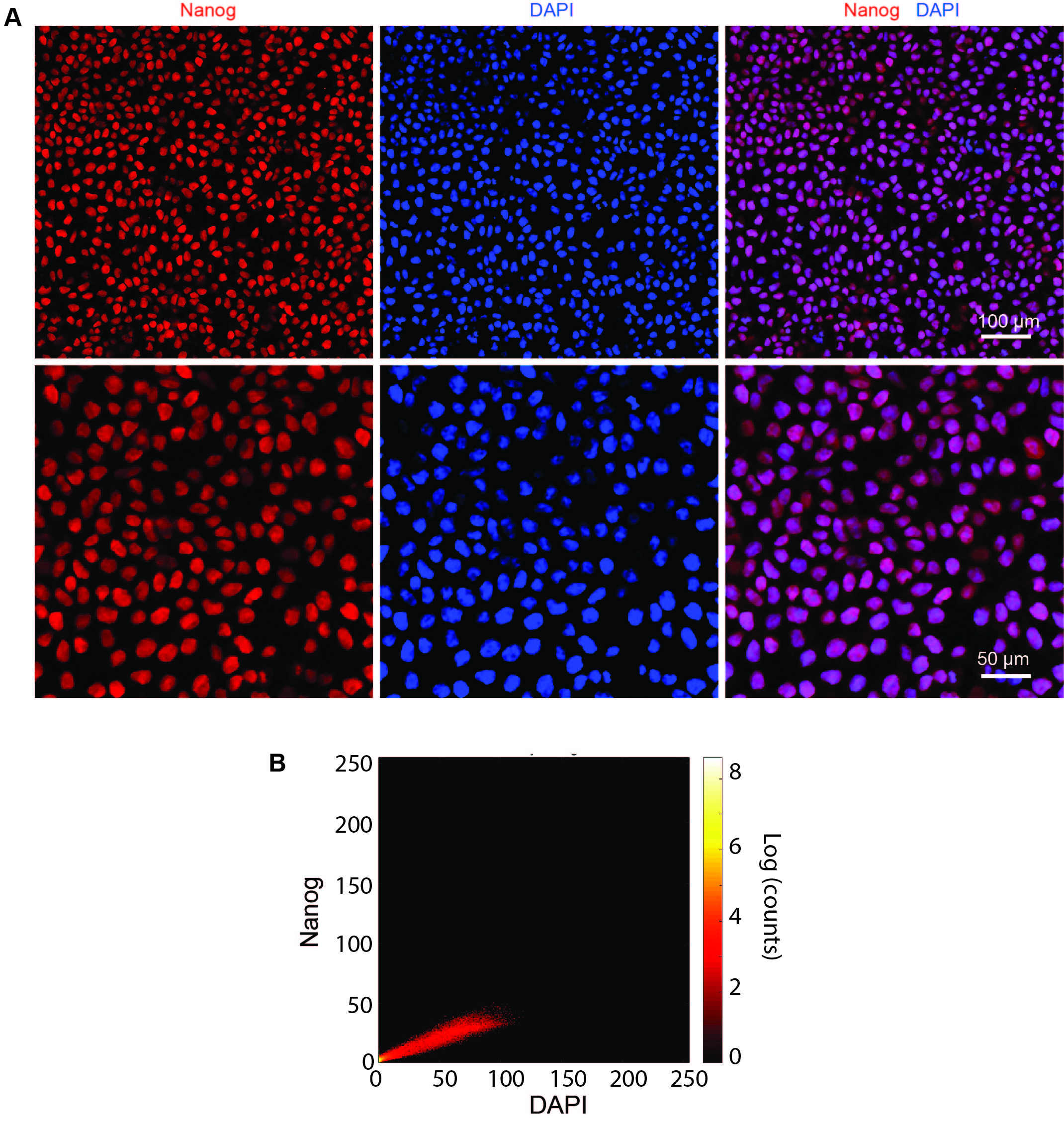

**Figure S5.** **Immunostaining images of hiPSCs in 2D culture confirm that cells are pluripotent stem cells.** (A) HiPSC cells are seeded in Matrigel coated 24-well glass bottom plate and cultured at 37 °C, 5% CO_2_ overnight. Then the cells are fixed with 4% paraformaldehyde (PFA) and stained with pluripotency marker, Nanog. Red and blue colors correspond to Nanog and DAPI, respectively. (B) 2D intensity histogram of Nanog and DAPI. The colocalization analysis is done by using Coloc2 Plugin in Fiji*.* Pearson coefficient is 0.97 between DAPI and Nanog. Costes statistical significance test is done by using 100 iterations, and the P value is 1.00.

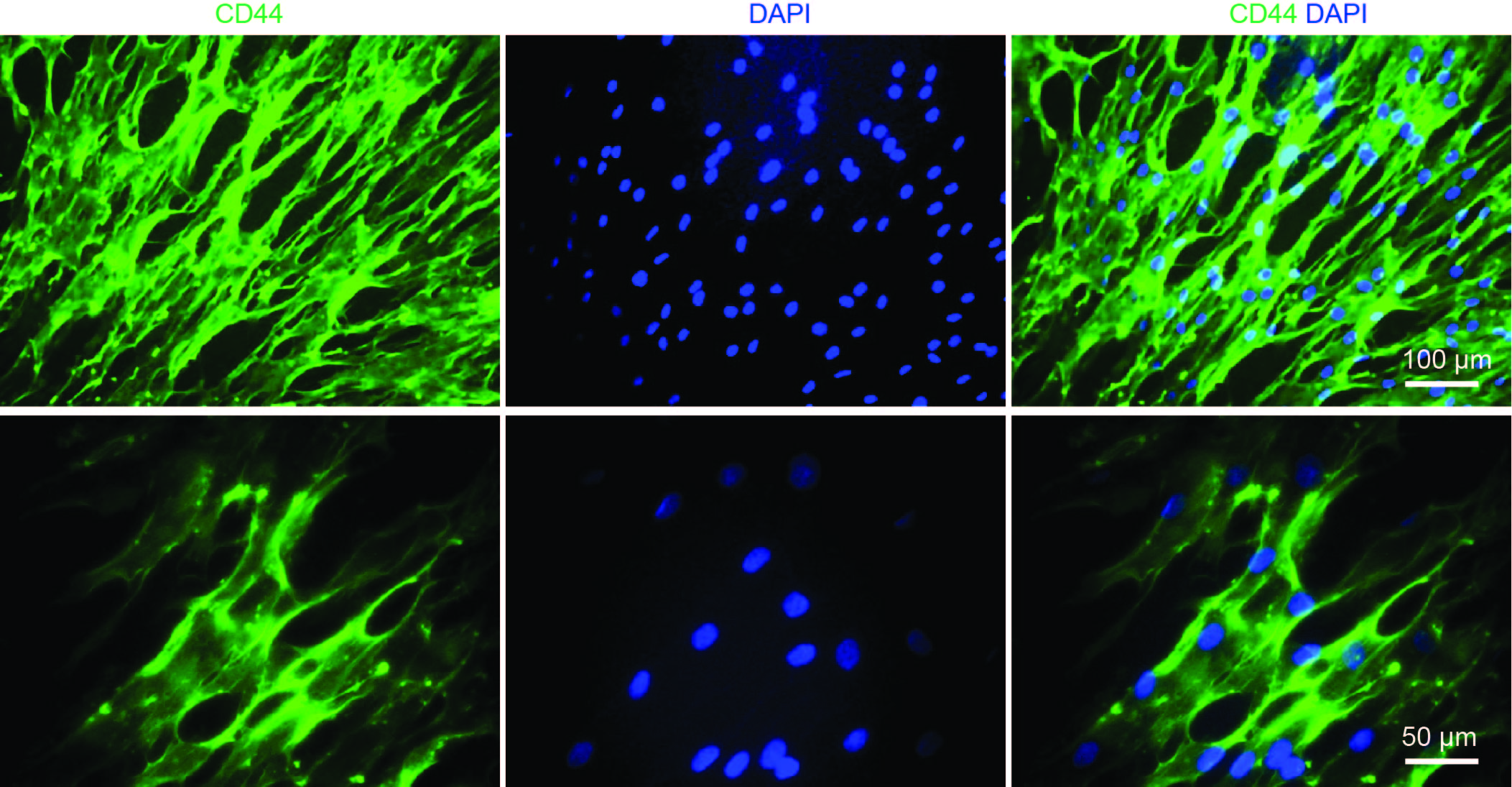

**Figure S6. Immunostaining images of 2D cultured hMSCs confirm the identity of hMSCs.** HMSCs are seeded in 24-well glass bottom plate and cultured at 37 °C, 5% CO_2_ overnight. Then the cells are fixed with 4% PFA and stained for protein marker CD44. Green and blue colors correspond to CD44 and DAPI, respectively.

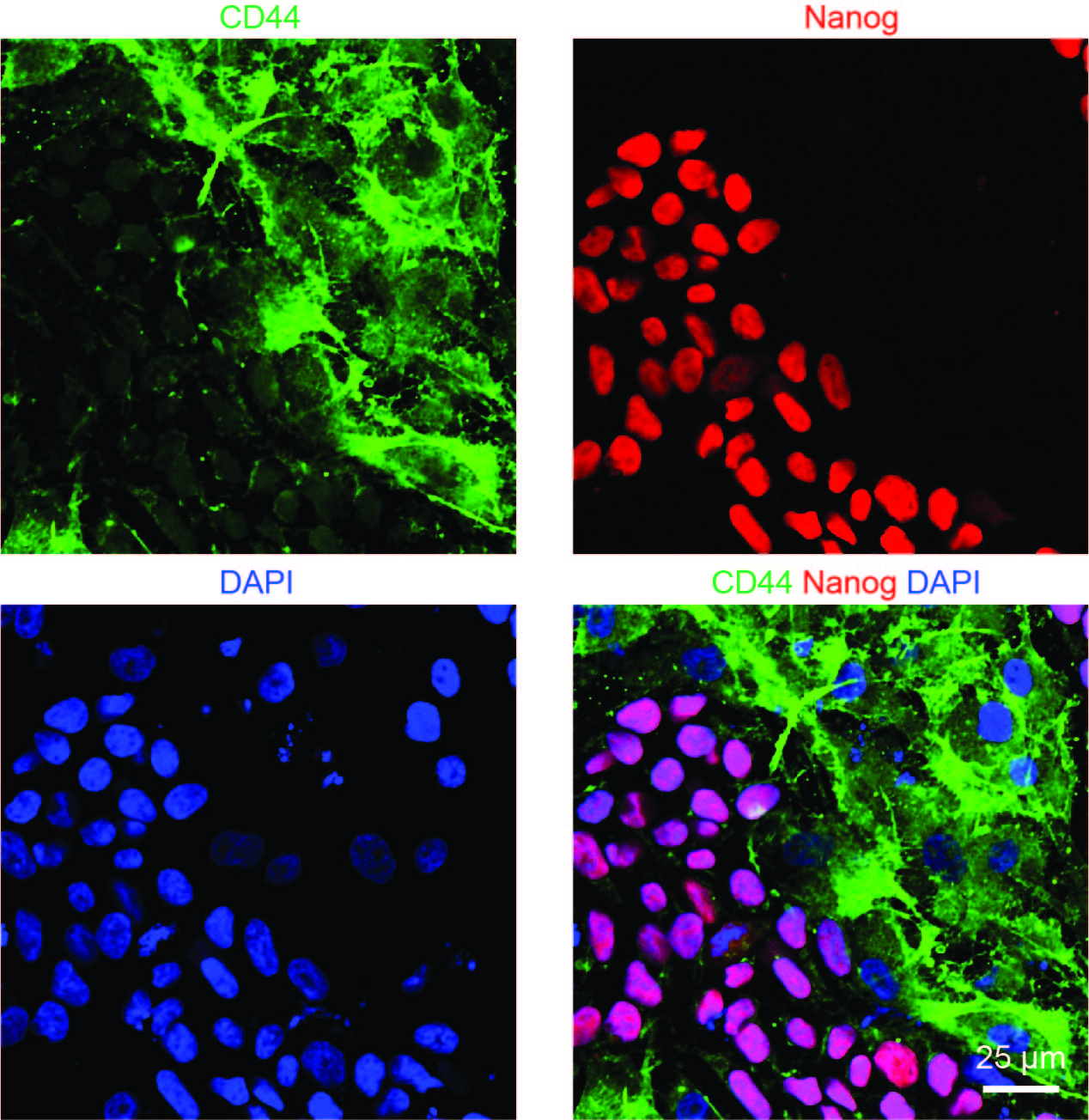

**Figure S7. Immunostaining images confirm the cell identities of co-cultured hMSCs and hiPSCs.** HiPSCs and hMSCs (1:1) are mixed and seeded in Matrigel coated 24-well glass bottom plate and cultured at 37 °C, 5% CO_2_ overnight. Then the cells are fixed with 4% PFA and stained for CD44 and Nanog. Green, red and blue colors correspond to CD44, Nanog, and DAPI, respectively.

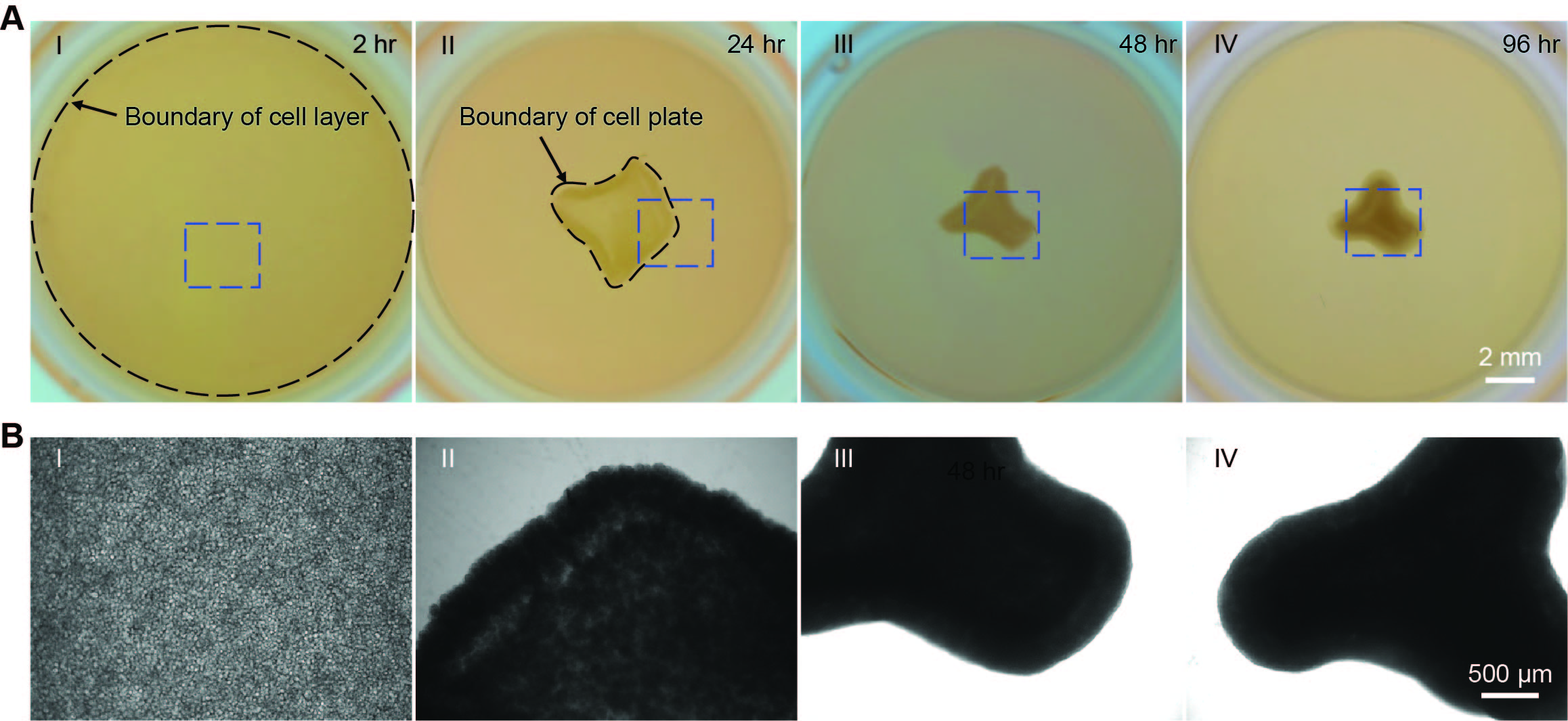

**Figure S8. HMSCs condensation drives 2D hMSCs/hiPSCs cell-sheet to fold into the 3D organoid.** HiPSCs (2-4×10^6^ cells) and hMSCs (2-3×10^5^ cells) are co-cultured on cured Matrigel in 24-well plate. The organoid is formed within 3-4 days. (A) Optical images show representative steps of organoid developmental process. Dark dash lines highlight boundaries of cell plate at different stages. (B) Zoom-in phase images of blue dashed boxes highlighted regions in (A) show the morphologies of cells or organoid at different stages.

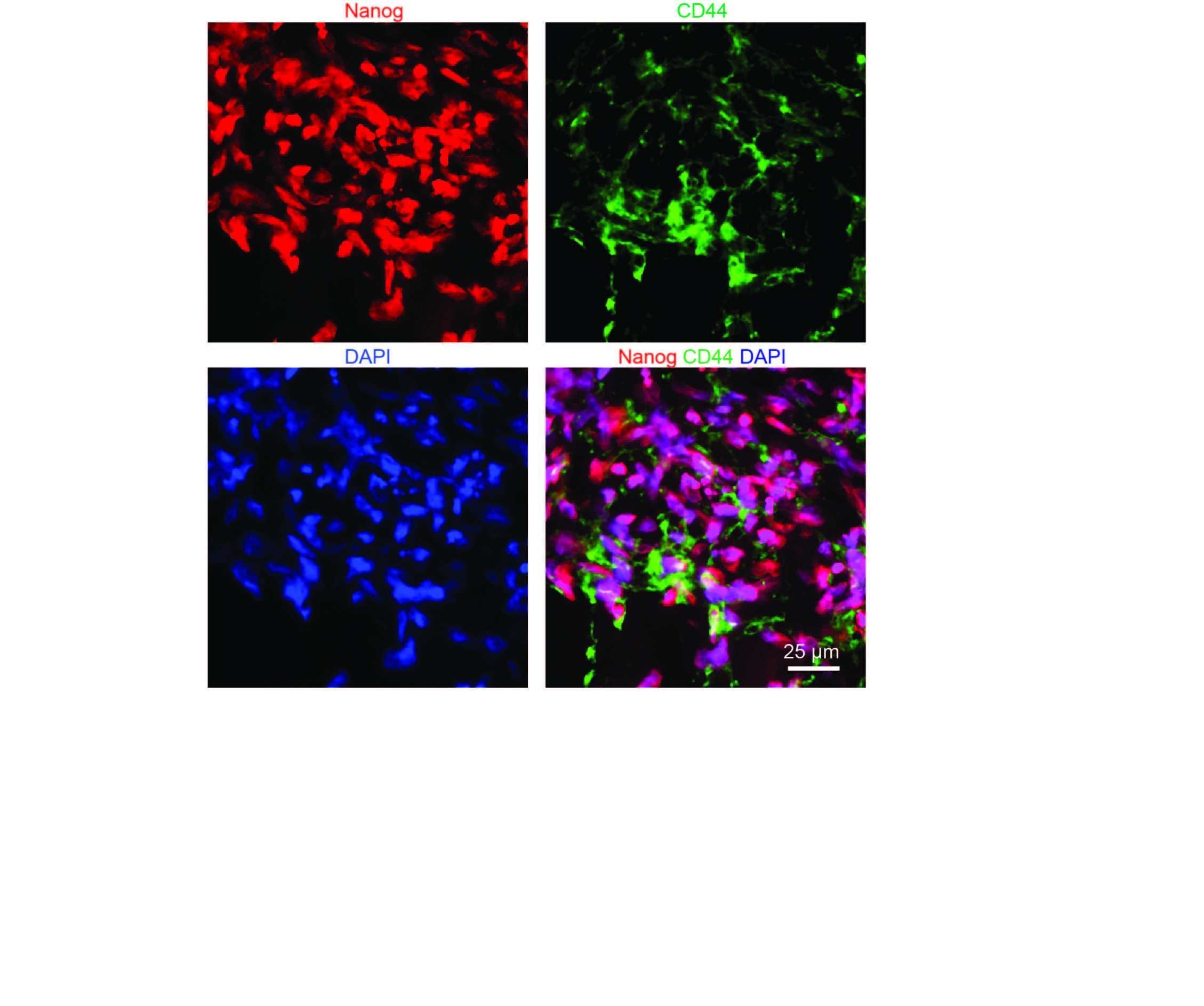

**Figure S9. Immunostaining images show the identities of hiPSCs and hMSCs in 3D organoid.** HiPSCs and hMSCs are co-cultured on cured Matrigel in 24-well plate to generate the organoid. The organoid was fixed, sectioned and stained after 4-day culture. Nanog and CD44 were immunostained and imaged for hiPSCs and hMSCs, respectively. Green, red and blue colors correspond to Nanog, CD44 and DAPI, respectively.

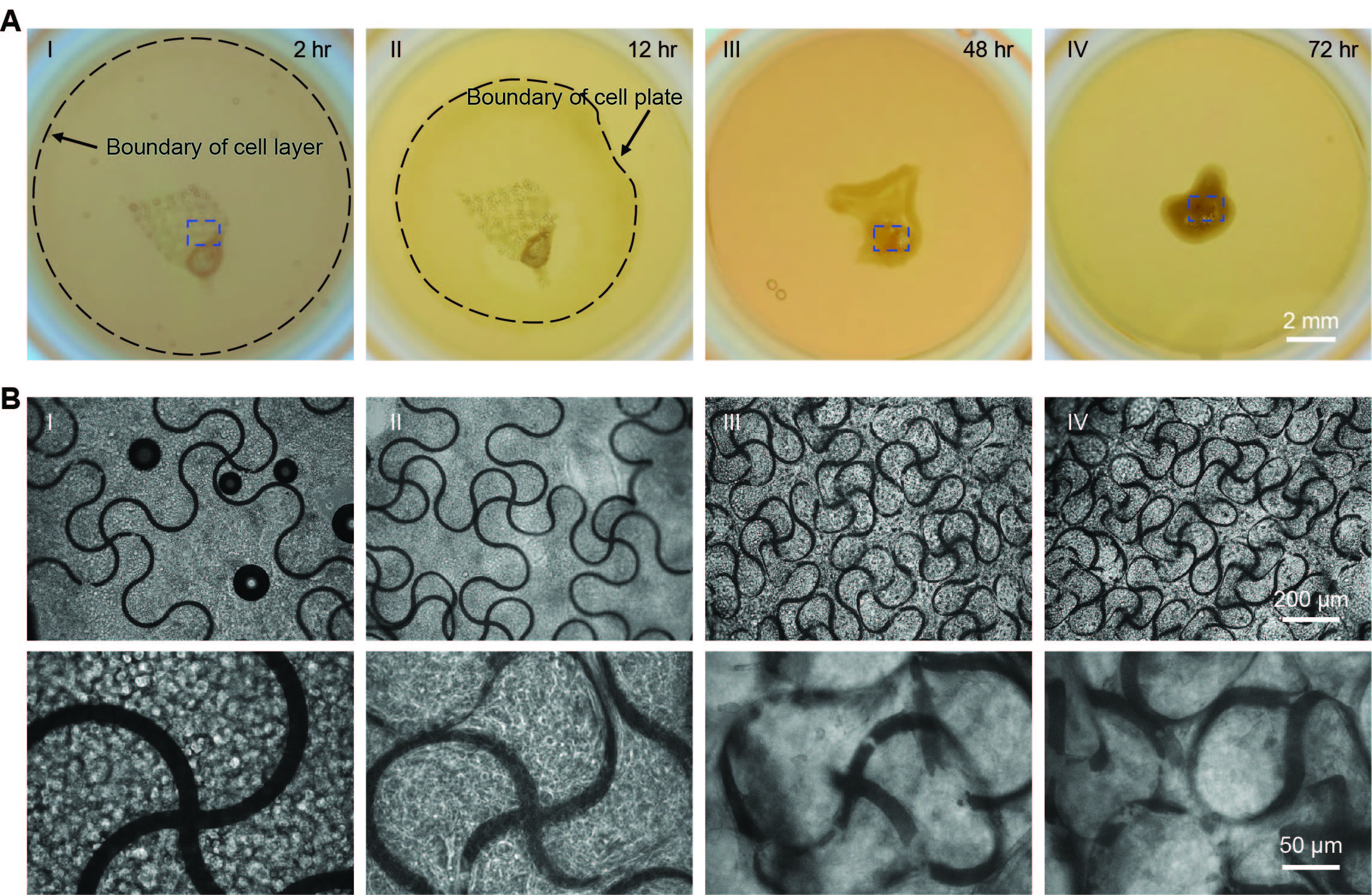

**Figure S10. HiPSCs/hMSCs organoid assembles stretchable mesh nanoelectronics with 20-μm-wide ribbons.** HiPSCs (2-4×10^6^ cells) and hMSCs (2-3×10^5^ cells) are co-cultured on cured Matrigel in 24-well plate with stretchable mesh nanoelectronics (*W/H =* 20 μm/0.8 μm). The cyborg organoid is formed within 3-4 days. (A) Optical images show representative steps of organoid organogenesis. Dark dash lines highlight boundaries of cell plate at different stages. (B) Zoom-in phase images show the structure evolution of stretchable mesh nanoelectronics inside organoids from blue dashed boxes highlighted regions in (A).

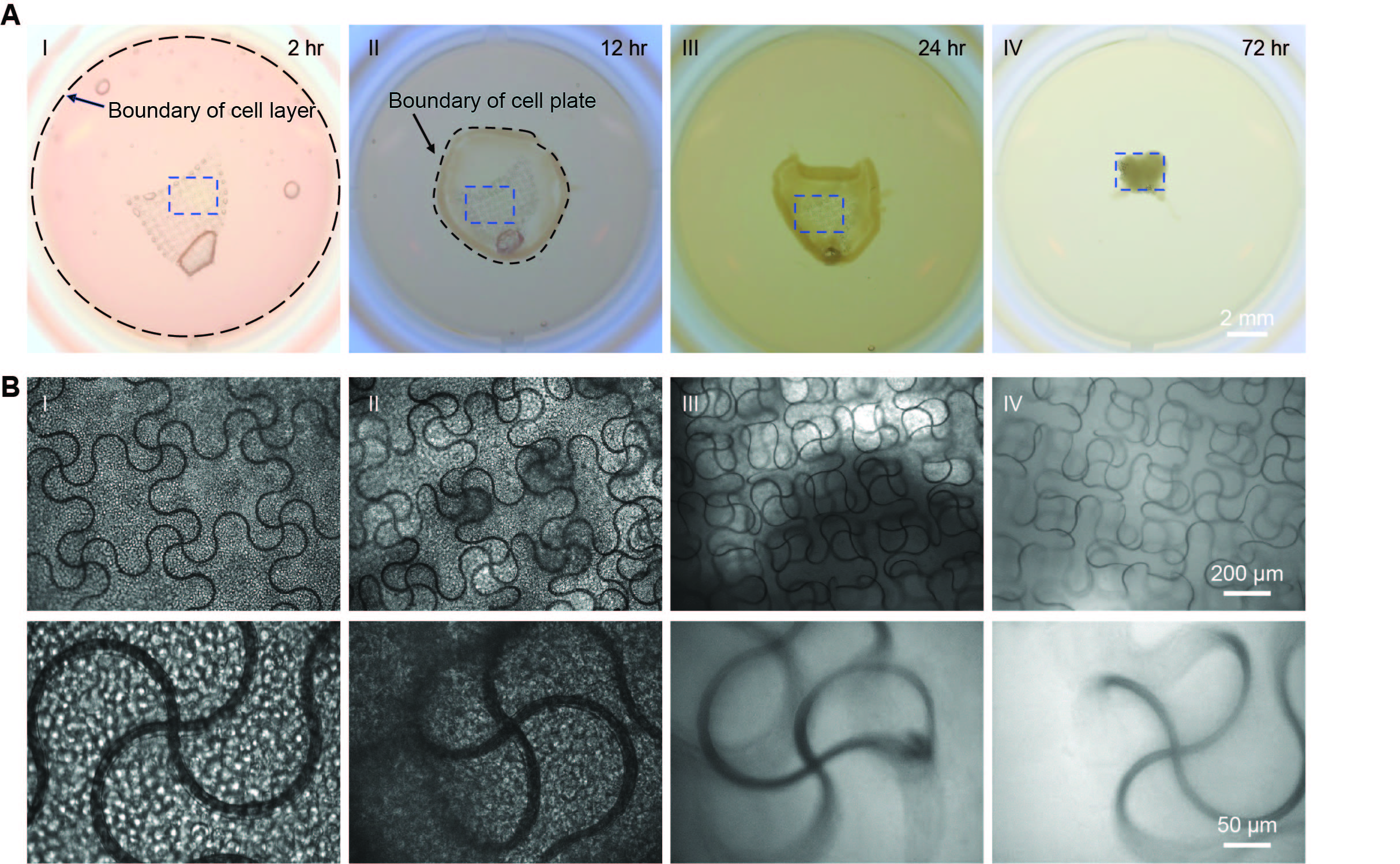

**Figure S11. HiPSCs/hMSCs organoid assembles stretchable mesh nanoelectronics with 10-μm-wide ribbons.** HiPSCs (2-4×10^6^ cells) and hMSCs (2-3×10^5^ cells) are co-cultured on cured Matrigel in 24-well plate with stretchable mesh nanoelectronics (*W/H =* 10 μm/2.8 μm). The cyborg organoid is formed within 3-4 days. (A) Optical images show representative steps of organoid organogenesis. Dark dash lines highlight boundaries of cell plate at different stages. (B) Zoom-in phase images show structure evolution of stretchable mesh nanoelectronics inside organoids from blue dashed boxes highlighted regions in (A).

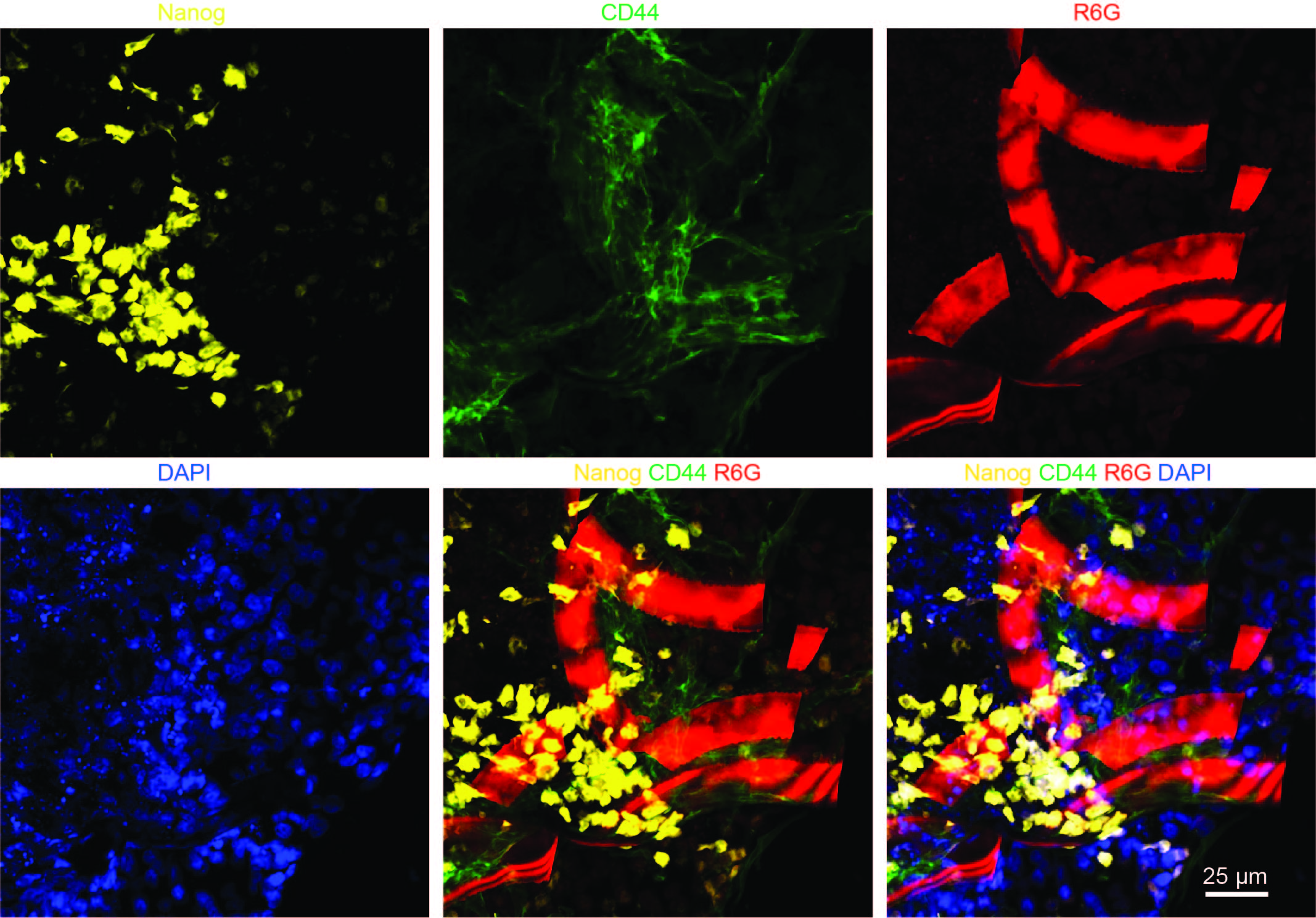

**Figure S12. Immunostaining images of hiPSCs/hMSCs cyborg organoid slices show the stretchable mesh nanoelectronics interwoven with hiPSCs and hMSCs.** HiPSCs (2-4×10^6^ cells) and hMSCs (2-3×10^5^ cells) are co-cultured on cured Matrigel in 24-well plate with stretchable mesh nanoelectronics (*W/H =* 20 μm/0.8 μm). The formed cyborg organoid is fixed, sectioned and stained. Fluorescence images of 30-μm thick organoid showed hiPSCs (labeled by Nanog) and hMSCs (labeled by CD44) in the organoid. R6G stained SU-8 structures from stretchable mesh nanoelectronics. Yellow, green, red and blue colors correspond to Nanog, CD44, R6G and DAPI, respectively.

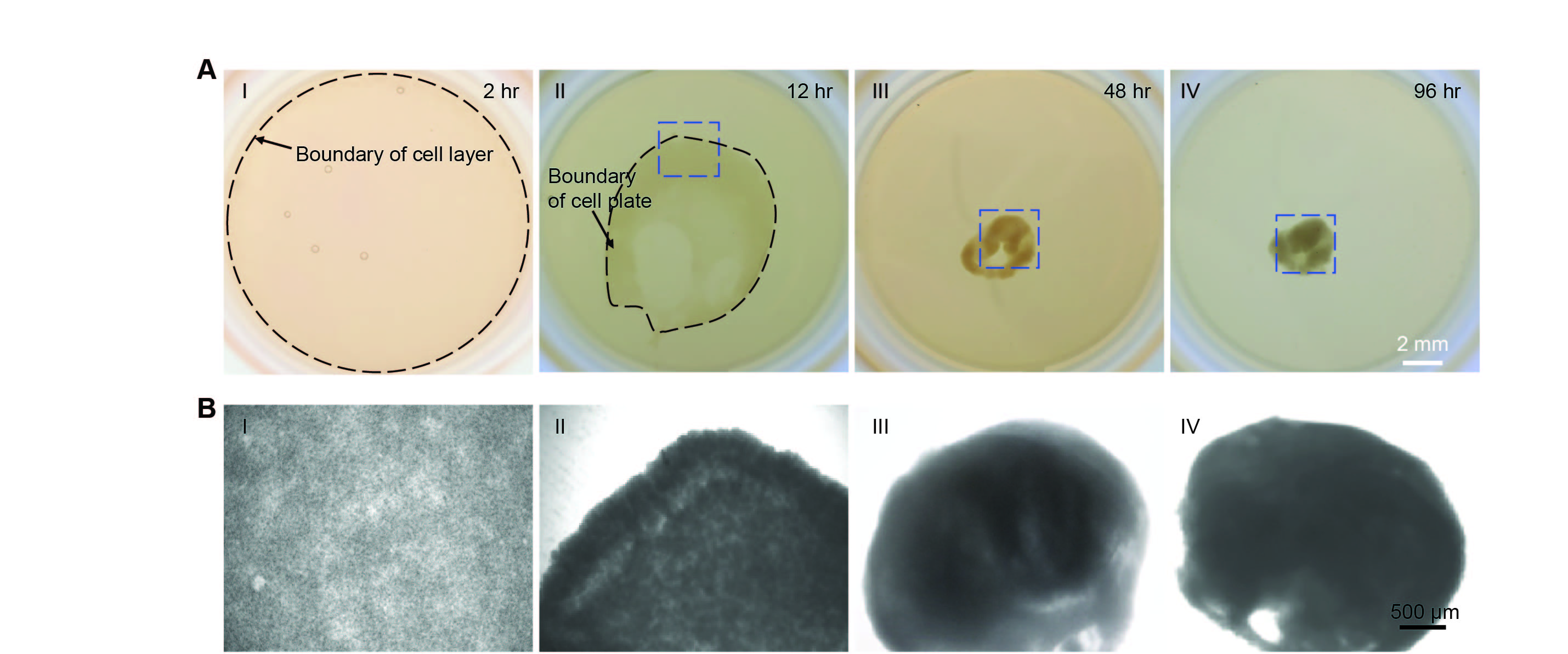

**Figure S13. HMSCs condensation drive 2D hMSCs/hiPSC-CPCs cell-sheet to fold into the 3D organoid.** HiPSC-CPCs (2-4×10^6^ cells) and hMSCs (2-3×10^5^ cells) are co-cultured on cured Matrigel in 24-well plate. The organoid is formed within 3-4 days. (A) Optical images show representative steps of organoid developmental process. Dark dash lines highlight boundaries of cell plate at different stages. (B) Zoom-in phase images show the morphologies of cells or organoid from blue dashed boxes highlighted regions in (A).

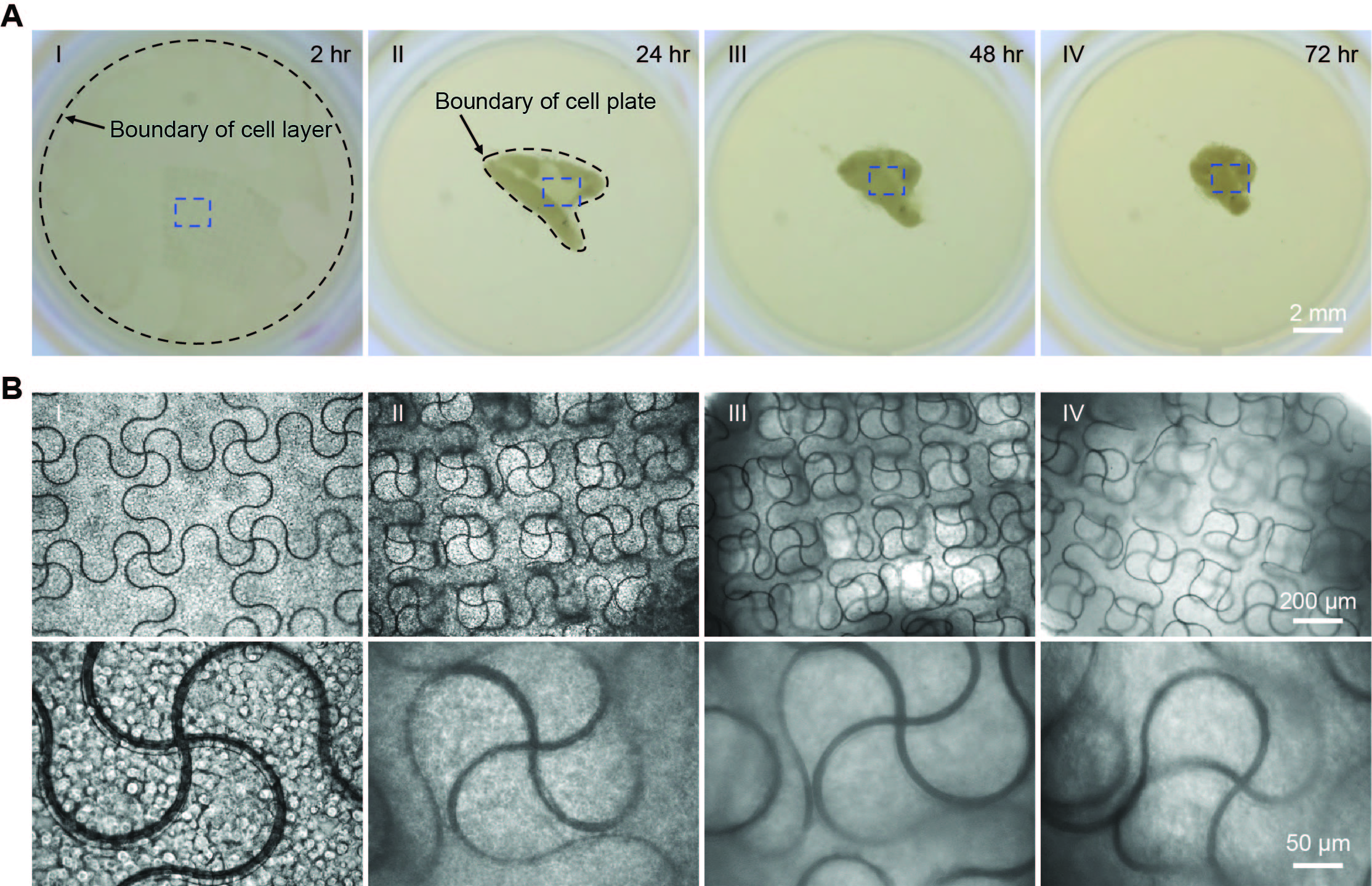

**Figure S14. hMSCs/hiPSC-CPCs organoid assembles stretchable mesh nanoelectronics with 10-μm-wide ribbons.** HiPSC-CPCs and hMSCs are co-cultured with stretchable mesh nanoelectronics (*W/H =* 10 μm/2.8 μm). The cyborg organoid is formed within 3-4 days. (A) Optical images show representative steps of organoid developmental process. Dark dash lines highlight boundaries of cell plate at different stages. (B) Zoom-in phase images show structure evolution of stretchable mesh nanoelectronics inside organoids from blue dashed boxes highlighted regions in (A).

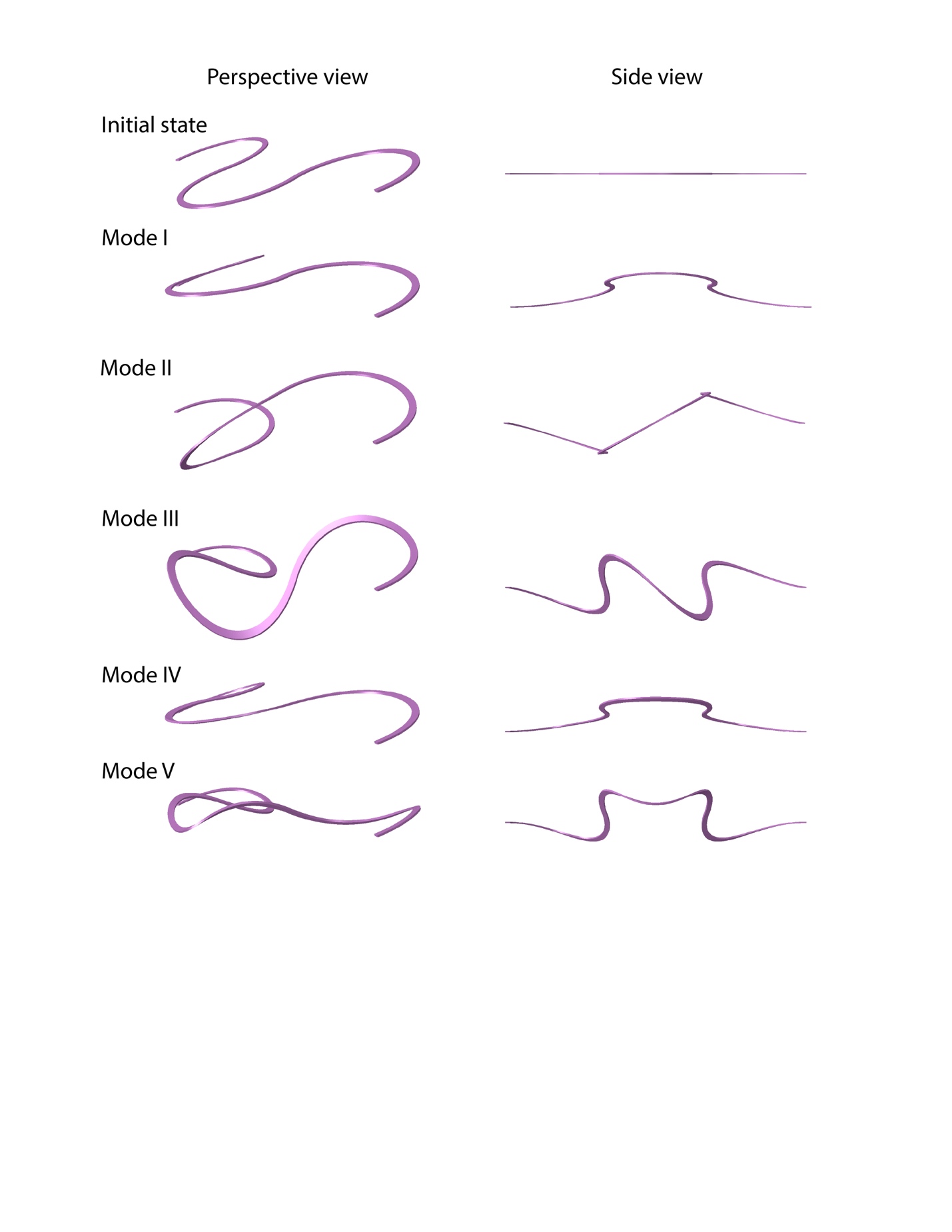

**Figure S15.** **Linear perturbation analysis shows buckling modes of a unit ribbon (*W/H =* 10 μm/2.8 μm) in compression.** Deformation in the Z direction is magnified 50 times for convenience.

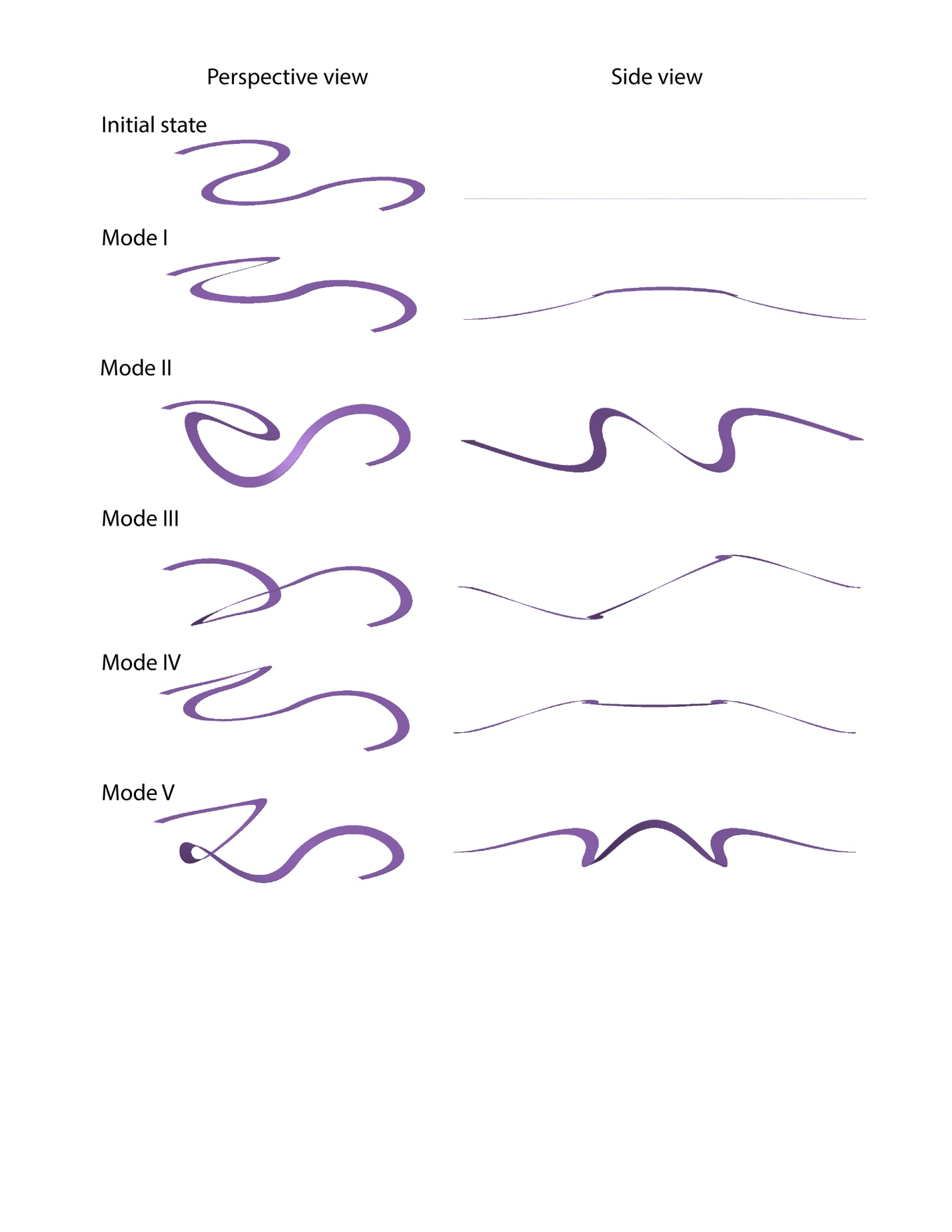

**Figure S16.** **Linear perturbation analysis shows buckling modes of a unit ribbon (*W/H =* 20 μm/0.8 μm) in compression.** Deformation in the Z direction is magnified 50 times for convenience.

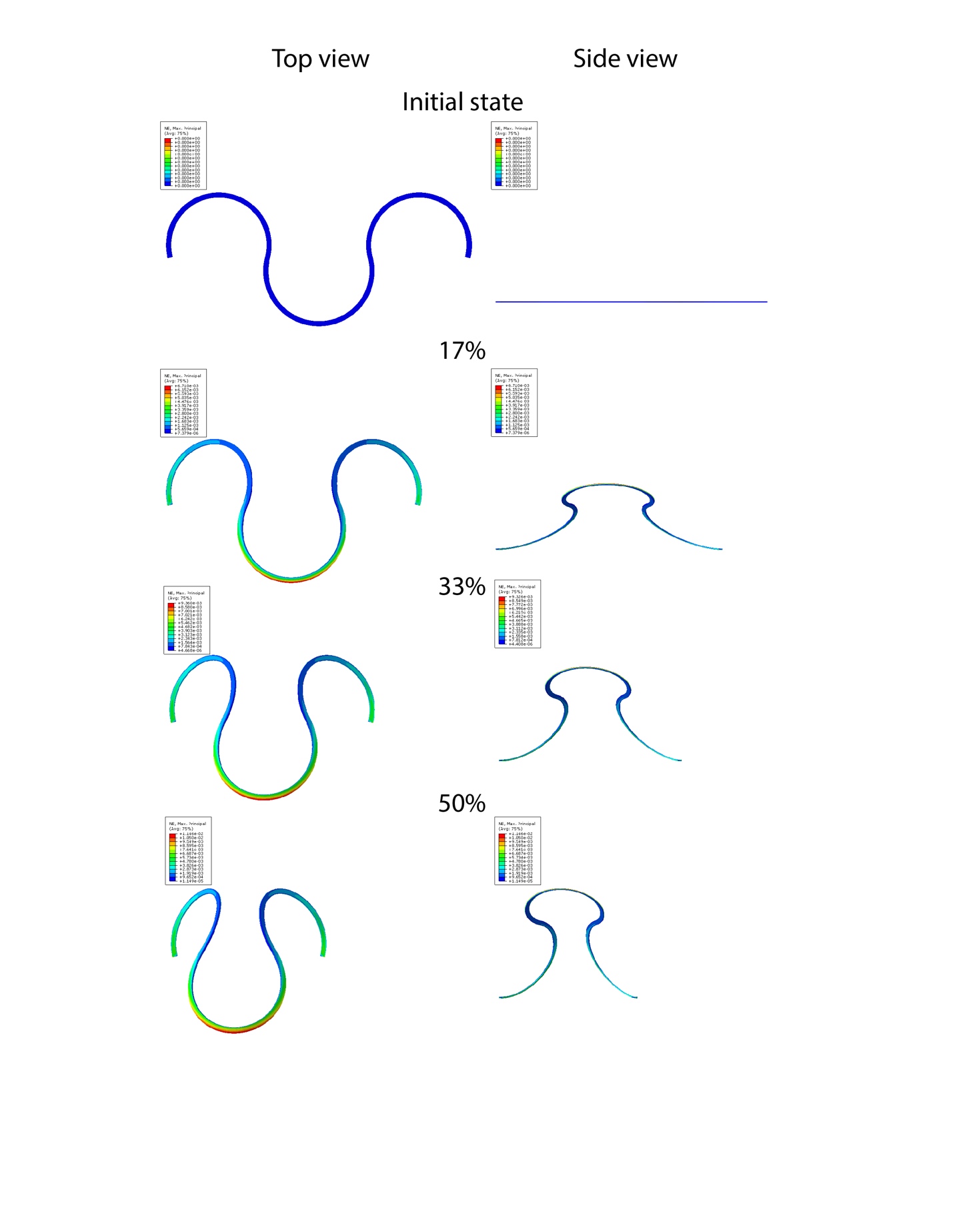

**Figure S17. FEA shows maximum principal nominal strain distribution during buckling (compression, Mode I) for a ribbon (*W/H =* 10 μm/2.8 μm), for an effective compressive strain from 0 to 50%.**

–––
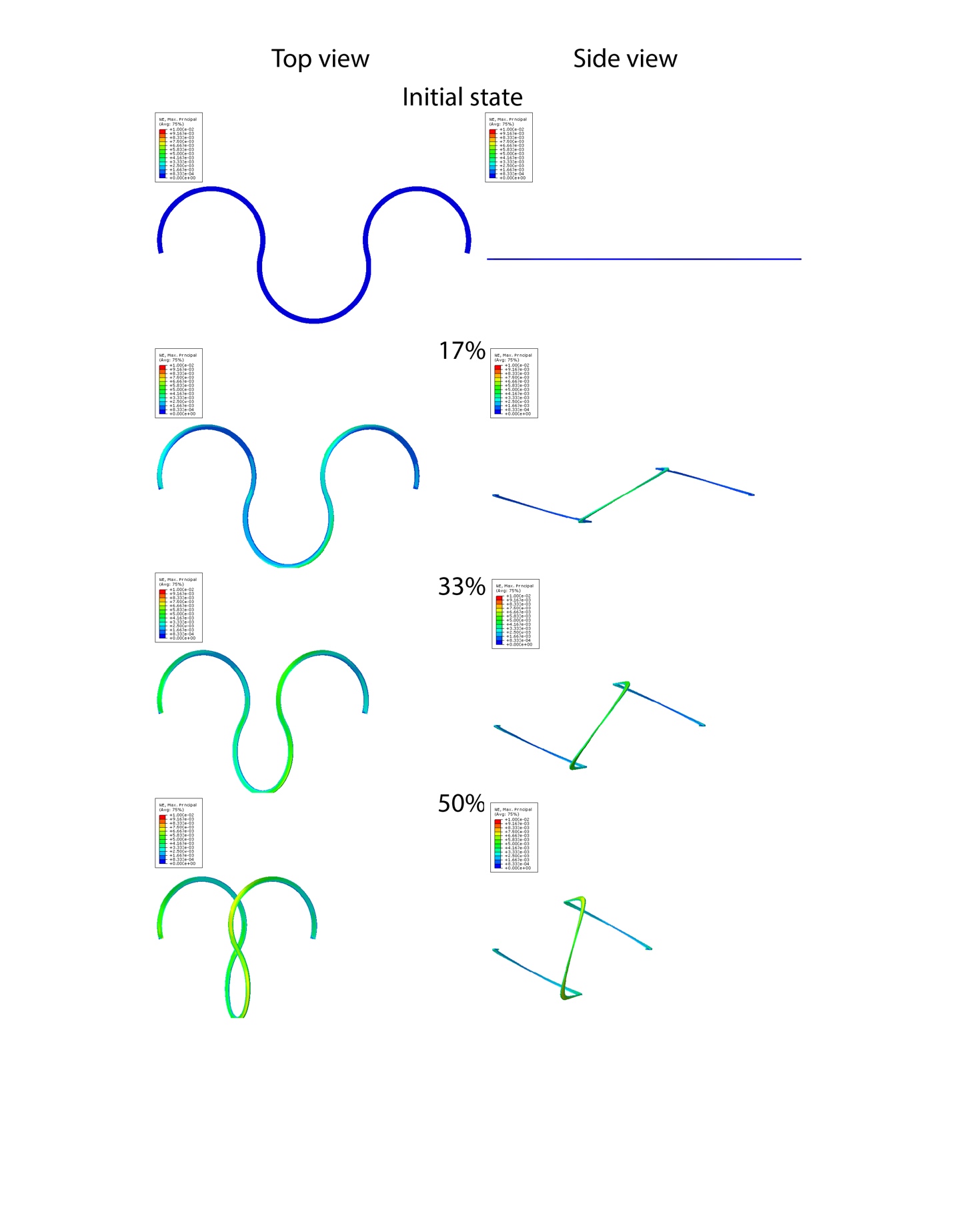

**Figure S18.** **FEA shows maximum principal nominal strain distribution during buckling (compression, Mode II) for a ribbon (*W/H =* 10 μm/2.8 μm), for an effective compressive strain from 0 to 50%.**

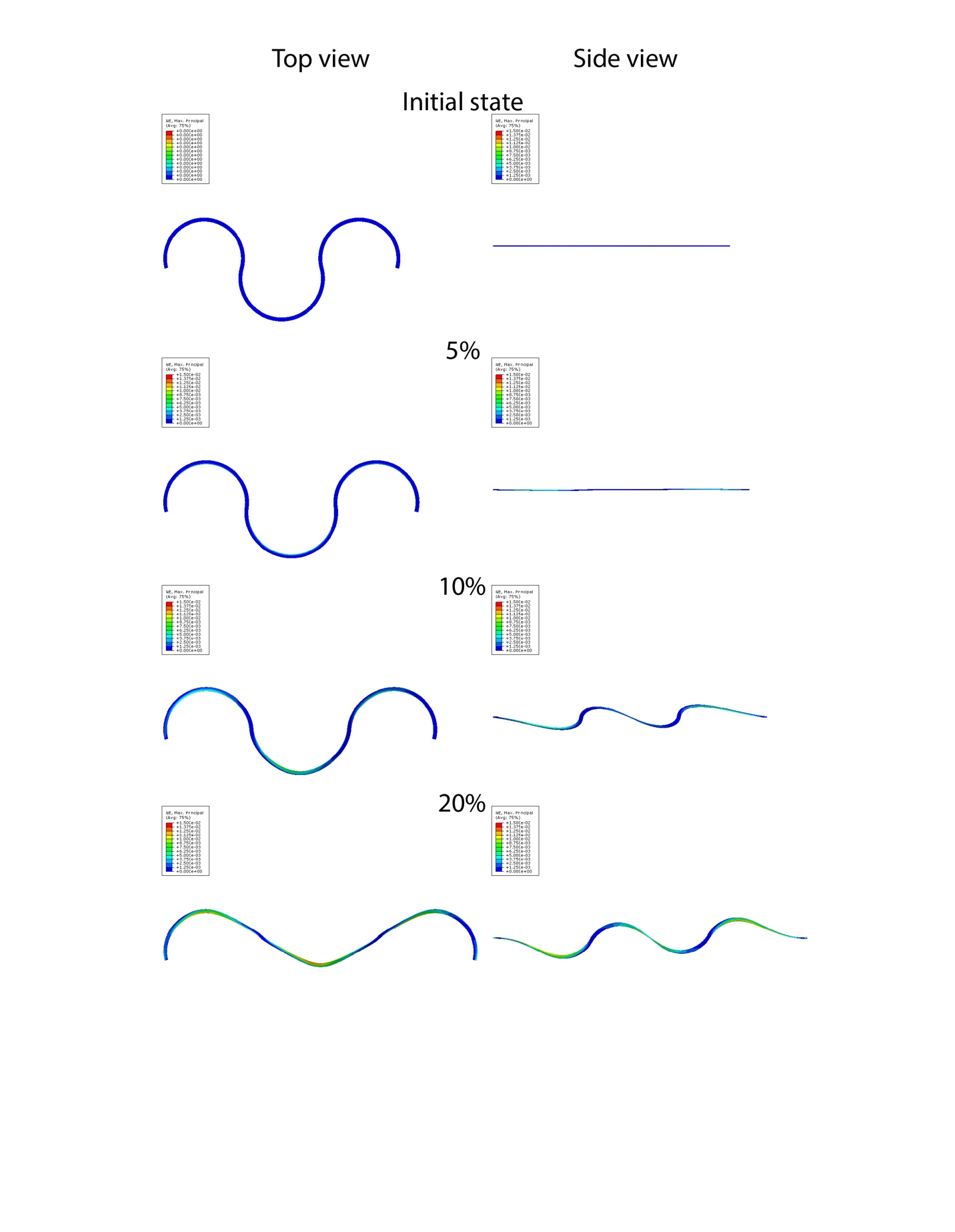

**Figure S19.** **FEA shows maximum principal nominal strain distribution during buckling (stretching, Mode I) for a ribbon (*W/H =* 10 μm/2.8 μm), for an effective tensile strain from 0 to 20%.**

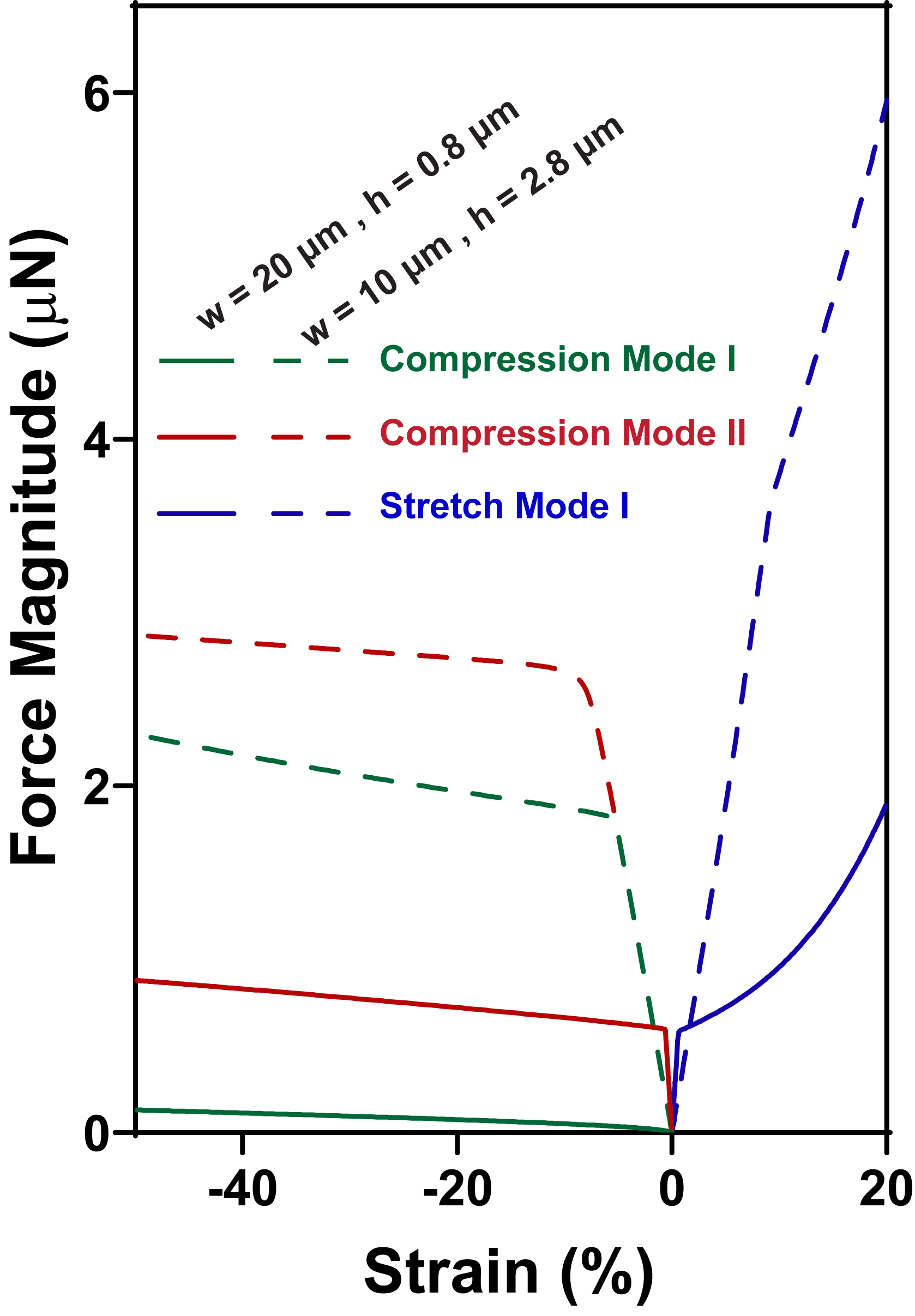

**Figure S20.** **Simulated result shows magnitude of the end-to-end reaction force on a unit ribbon as a function of the applied effective in-plane strain.** The force magnitude is higher for the thicker device as expected. The bend in each curve corresponds to the onset of buckling in the simulations.

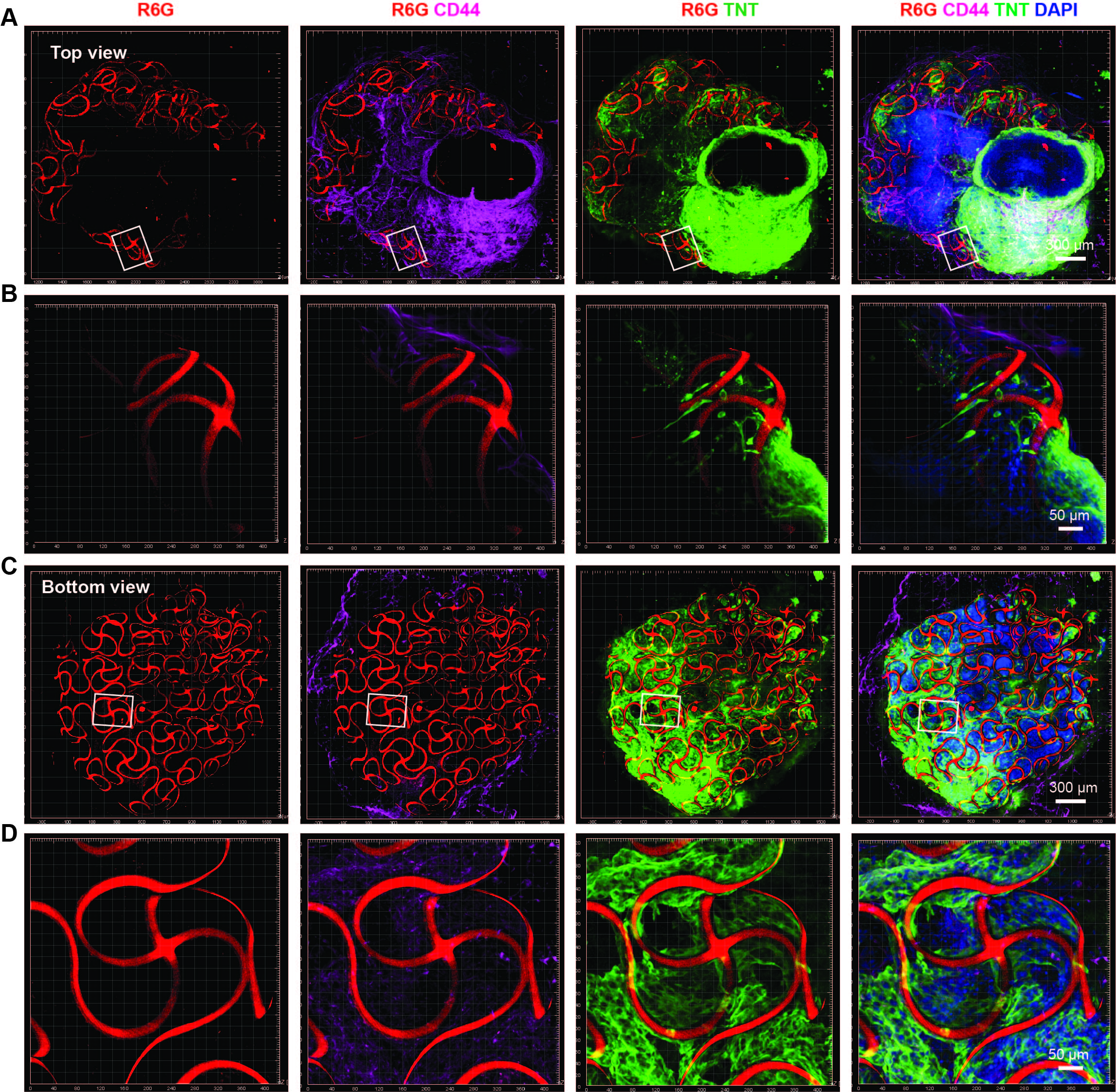

**Figure S21. 3D reconstructed confocal images from cleared cyborg organoids show the 3D structure of stretchable mesh nanoelectronics and its integration with cells.** The cyborg cardiac organoid is fixed, cleared and immunostained. Top (A) and bottom (C) views of fluorescence imaging show 3D structures of the stretchable mesh nanoelectronics in the cyborg cardiac organoid. (B, D) Zoom-in views from the white boxes highlighted region in (A, B) show coupling between cardiomyocytes or hMSCs with nanoelectronics. Green, red, magenta and blue colors correspond to TNT, CD44, R6G and DAPI, respectively.

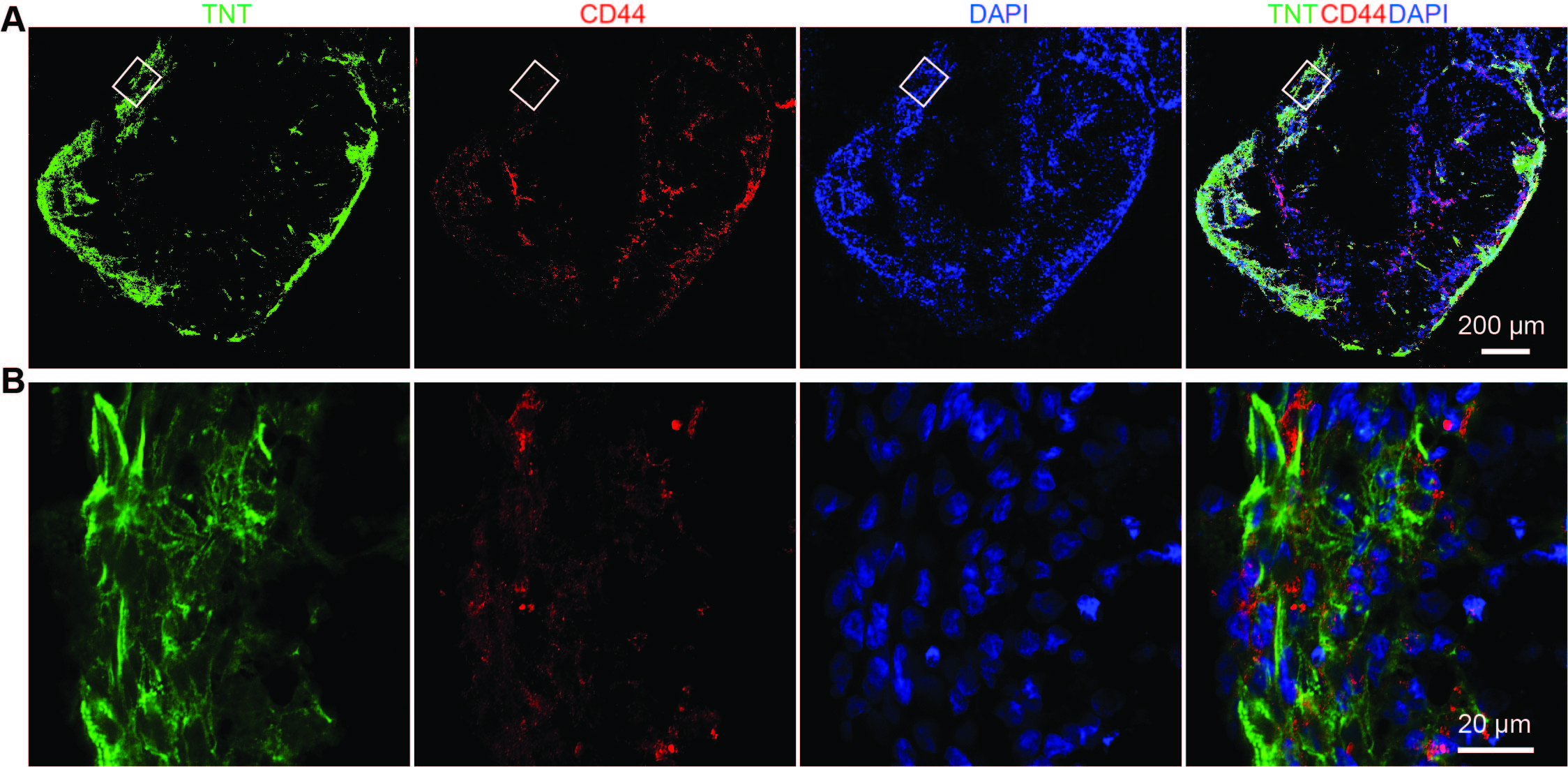

**Figure S22. Immunostaning images show differentiated hiPSCs-derived cardiomyocytes and hMSCs in cardiac organoids.** HiPSC-CPCs and hMSCs are co-cultured. The formed cardiac organoid is fixed, sectioned and imaged after 40 days differentiation. (**A**) Projection of 3D reconstructed confocal microscopic fluorescence images of 30-μm thick organoid showed cardiomyocytes (labeled by TNT) and hMSCs (labeled by CD44) in the organoid. (**B**) Zoom-in views show sarcomere alignment in the organoid from the white box highlighted in (**A**). Green, red and blue colors correspond to TNT, CD44, and DAPI respectively.

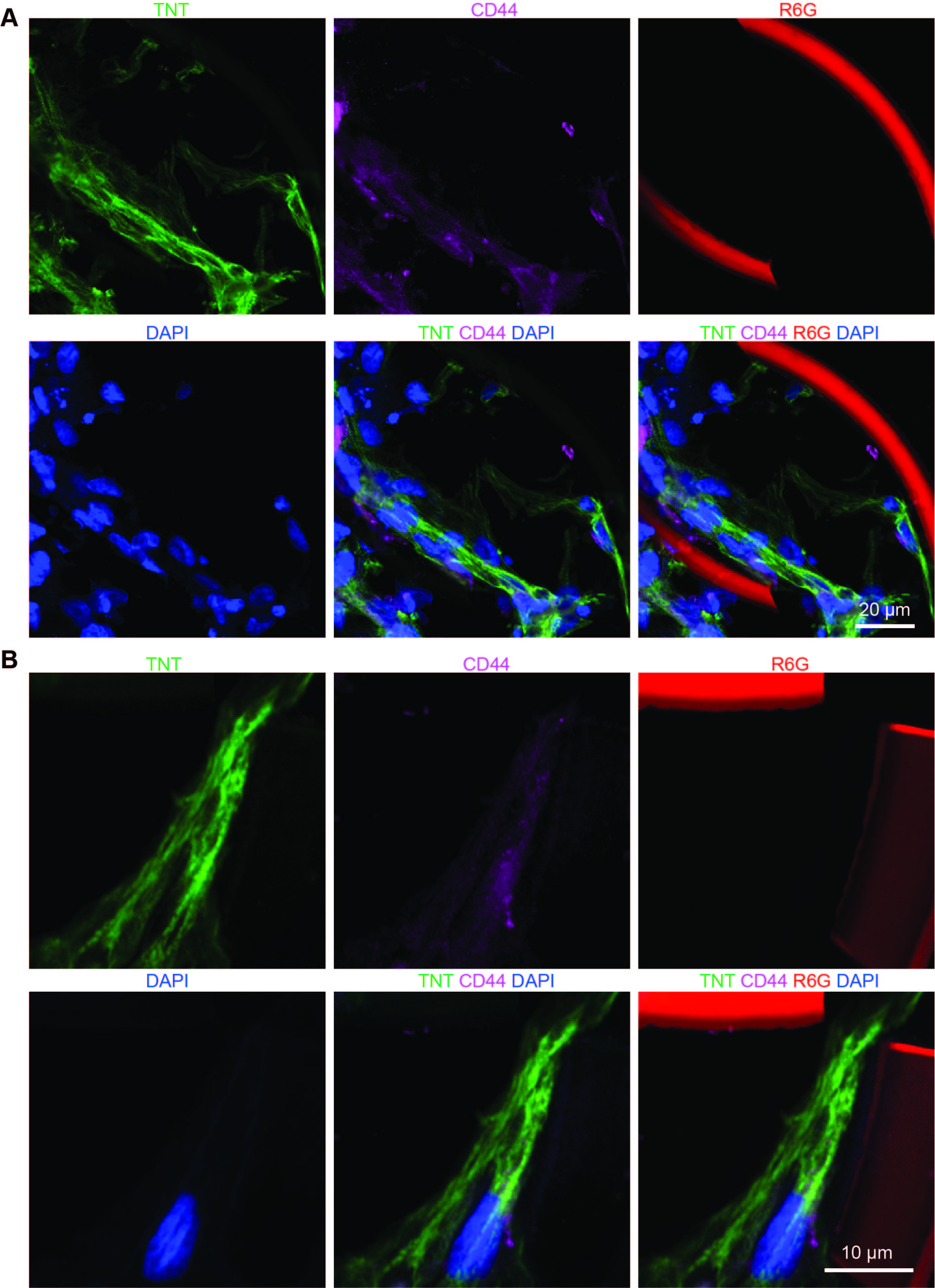

**Figure S23. Immunostaining images show differentiated hiPSCs-derived cardiomyocytes and hMSCs in cardiac cyborg organoids.** HiPSC-CPCs and hMSCs are co-cultured with stretchable mesh nanoelectronics (*W/H =* 10 μm/2.8 μm). The formed cyborg cardiac organoid is fixed, sectioned and imaged after 40 days differentiation. (**A**) Confocal microscopic fluorescence images of 30-μm thick organoid show TNT+ cardiomyocytes and CD44+ hMSCs in the organoid. (**B**) Zoom-in views show subcellular localization of TNT and CD44 in the organoid. Green, magenta, red and blue colors correspond to TNT, CD44, R6G and DAPI respectively.

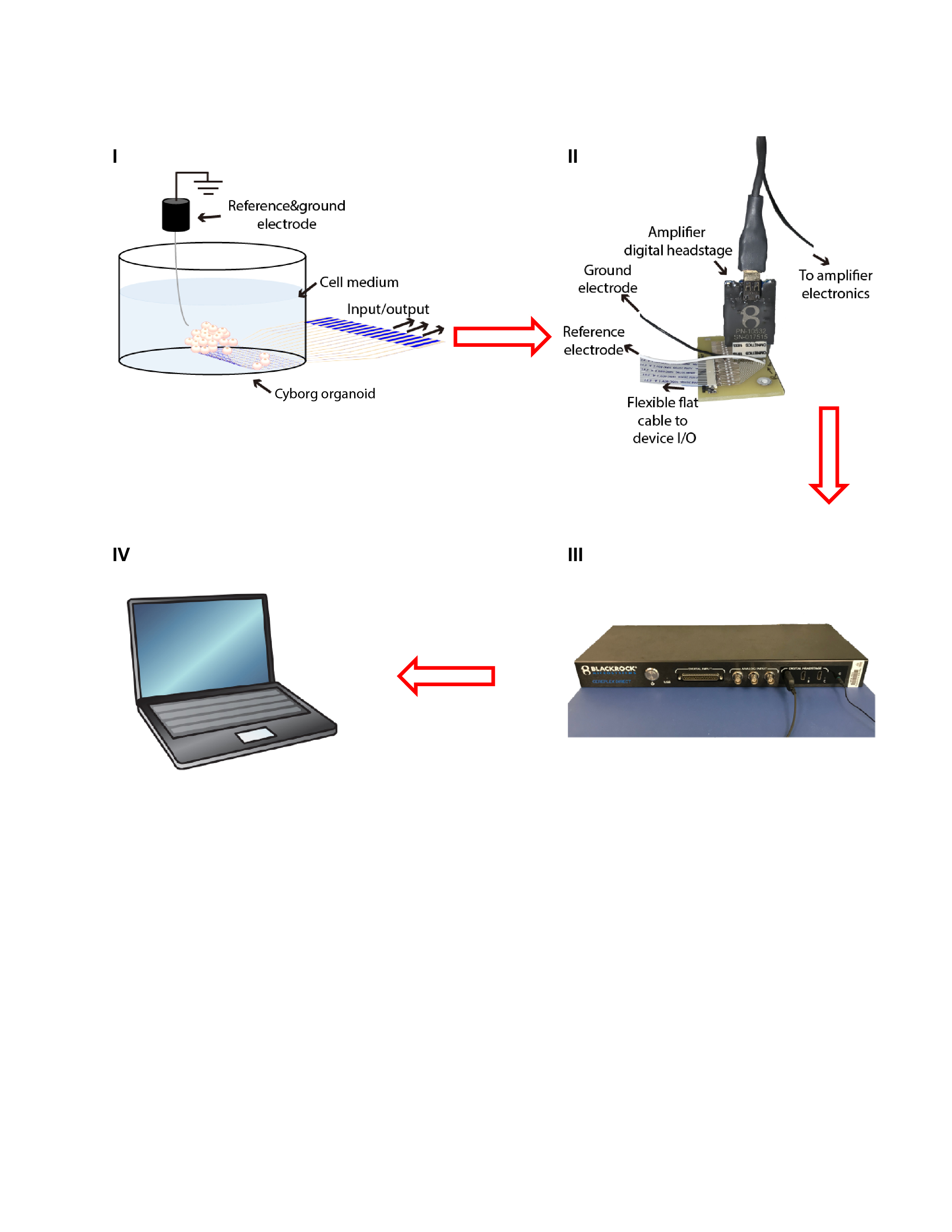

**Figure S24.** **Schematics of the electrophysiological measurement setup.** (I) The front end of the device (cyborg organoid) that is released in an enclosed chamber containing reference and ground electrodes, and the rare end of the device (input/output) that is soldered with a 16-pin flexible flat cable (FFC). (II) A customized printed circuit board that joins the 16-pin FFC connector and the Blackrock μ digital headstage together. (III) The digital headstage is connected to the amplifier electronics (Blackrock CerePlex Direct voltage amplifier), which feeds the signals through USB into a computer (IV) for processing.

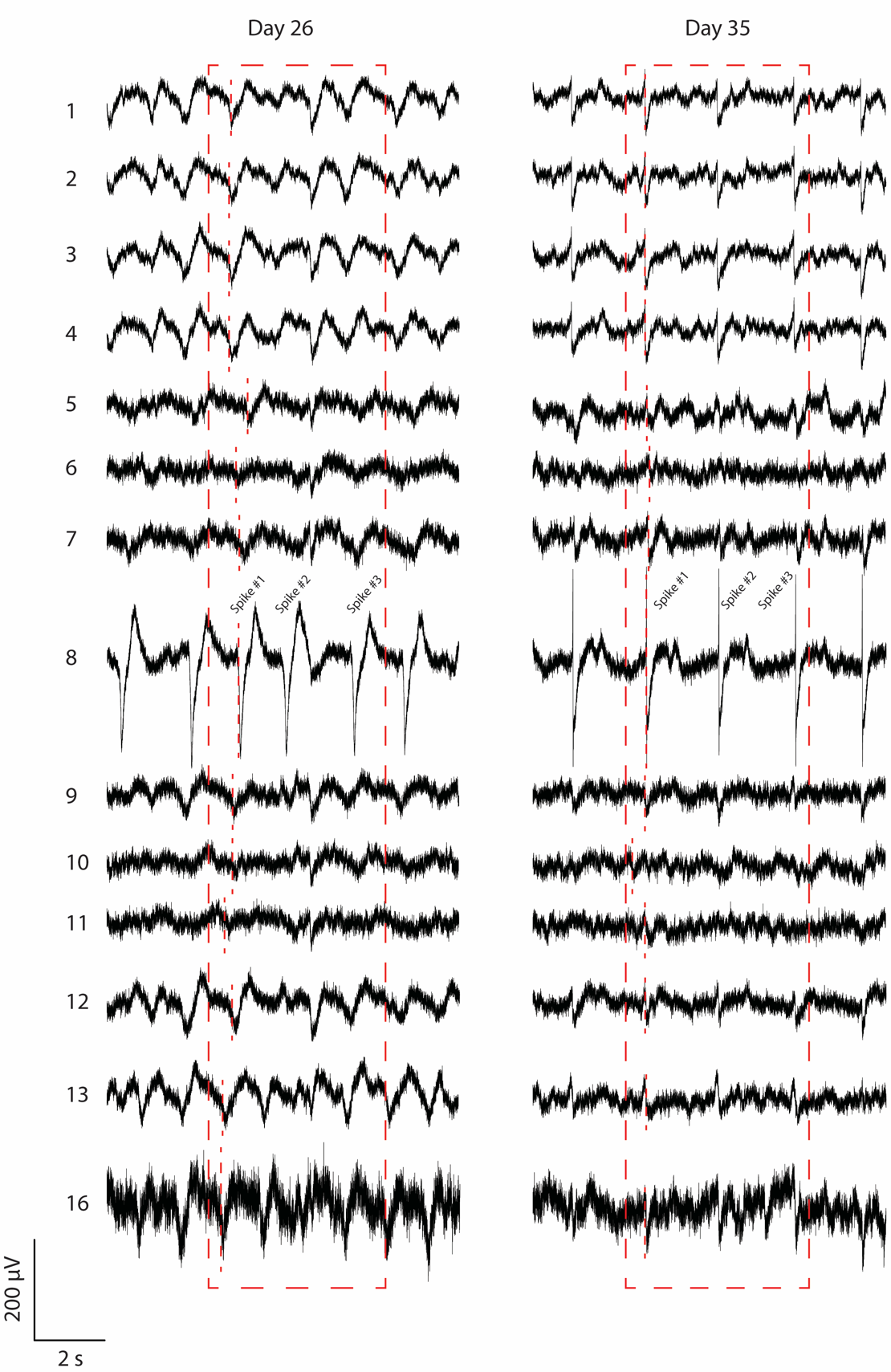

**Figure S25.** **Electrophysiology data for 14 channels during day 26 and 35 of culture shows transition from irregular activity patterns to synchronous beating.** This dataset was used to build the activation maps in Figure 4H, for three consecutive spiking events inside the dashed red rectangles. Red dashed segments at day 26 represent the activation time for spike #1 for each channel.

Movie S1.

Folding of the released device in solution.

**Movie S2.**

Stretching of the released device in solution.

**Movie S3.**

4X magnification phase imaging video shows the beating cardiac cyborg organoid at day 30 of differentiation.

**Movie S4.**

Zoom-in video (10X) shows the beating cardiac cyborg organoid at day 30 of differentiation.

**Movie S5.**

Further zoom-in video (40X) shows the beating cardiac cyborg organoid at day 30 of differentiation.

**Movie S6.**

Control: the beating cardiac organoid at day 30 of differentiation.
